# Dark CO_2_ Fixation via the Ethylmalonyl-CoA Pathway Establishes Metabolic Parity: A Stoichiometric Basis for Compounding Ecosystem Shifts

**DOI:** 10.64898/2026.08.26.747204

**Authors:** Yubo Wang, Scott van der Veer, Timothy Paez Watson, Martin Pabst, Dimitry Y. Sorokin, Mark C.M. van Loosdrecht

## Abstract

A central challenge in microbial ecology is understanding how competing guilds coexist and why community structures unexpectedly drift away from theoretical steady states. While polyphosphate accumulation provides energy and redox buffering in many heterotrophs, how competing lineages occupying the same ecological niche without polyphosphate synthesis manage equivalent intracellular redox imbalances remains unposed. Using an enhanced biological phosphorus removal macrocosm, we integrated quantitative stoichiometry with metaproteomics to resolve alternative metabolic strategies that underpin cellular homeostasis in heterotrophs. We show that glycogen-accumulating organisms (GAOs) achieve baseline metabolic parity with polyphosphate-accumulating organisms (PAOs) through a parallel redox-buffering mechanism: heterotrophic CO_2_/HCO_3_^−^ re-assimilation via the ethylmalonyl-CoA pathway. This inorganic carbon fixation couples structural carbon conservation with tight redox control, mitigating intracellular electron overflow and eliminating the long-assumed GAO bioenergetic inferiority. The bioenergetic efficiency of GAOs is further fortified by fine-tuned metabolic wiring, featuring energy-efficient high affinity acetate activation, ferredoxin-centered biochemistry, and energy-neutral polyhydroxyalkanoates (PHA) mobilization. Strikingly, minor formate co-feeding disrupted this established PAO–GAO parity. Stoichiometric simulations revealed that this formate supplementation creates an asymmetric bioenergetic niche that grants per-cycle energy gains exclusively to GAOs. Decoupled from hydraulic throughput, solids retention time control retains cells carrying accumulated intracellular inventory, translating subtle per-cycle stoichiometric edges into a multi-generational ratchet and drives a rapid community shift from PAO–GAO co-dominance to GAO dominance. By exposing limitations of traditional single-substrate and single-cycle steady-state models, our findings reveal how inorganic resource management and generational metabolic compounding govern community assembly in biomass-retaining microbial ecosystems.

## Introduction

Maintaining intracellular homeostasis is essential for all cellular life. In response to metabolic stresses arising from imbalances in redox state (NAD(P)H/NADP^+^), energy charge (ATP/ADP), intracellular osmolarity, or localized chemical toxicity, many prokaryotes accumulate specialized intracellular storage polymers, including polyhydroxyalkanoates (PHAs), glycogen, cyanophycin, polyphosphate and polysulfur. Rather than functioning merely as passive nutrient stockpiles, these macromolecules serve as dynamic physiological buffers tailored to specific environmental pressures. Specifically, PHAs and glycogen primarily mitigate carbon overflow and redox imbalances^1,2^, cyanophycin sequesters excess amino acids to alleviate osmotic stress^3,4^, and polyphosphate serves as a versatile energy and phosphate buffer^5,6^. In sulfide-rich environments, polysulfur accumulation operates as a kinetic detoxification strategy and simultaneously serves as a respiratory electron reserve^7^. These diverse adaptive traits are particularly pronounced in ecosystems characterized by fluctuating electron donor/acceptor pools and dynamic nutrient availability, such as the estuarine habitats^8^. Several of these storage mechanisms, most notably those governing polyphosphate, PHA, and glycogen dynamics, have been successfully exploited in enhanced biological phosphorus removal (EBPR) systems, representing one of the most widely implemented environmental biotechnologies for global wastewater treatment^9^.

Beyond polymer accumulation, sustaining cellular homeostasis under dynamic conditions requires a highly coordinated interface between catabolism and anabolism. To balance dynamic flux branches and prevent metabolic bottlenecks, prokaryotes rely on central carbon networks that fulfill critical anaplerotic functions; among these, the glyoxylate shunt is the most well-known mechanism for acetyl-CoA assimilation^10,11^. Another prominent route for acetyl-CoA assimilation is the ethylmalonyl-CoA (EMC) pathway, which operates at the core of carbon metabolism in diverse Alphaproteobacteria^12–20^ (e.g., *Rhodobacter sphaeroides*^12^ and *Methylobacterium extorquens*^13–16^*, Methylosinus trichosporium OB3b*, and *Methylocystis sp. MJC1*^17–20^), as well as select Actinomycetota lineages (e.g., *Streptomyces* spp.^21^). While the EMC pathway’s primary structural role is to convert acetyl-CoA into essential precursor metabolites for biomass synthesis, such as pyruvate, phosphoenolpyruvate, oxaloacetate and α-ketoglutarate^22^, its unique architecture presents distinct physiological features when compared to the carbon-neutral glyoxylate shunt. Notably, via its integrated reductive CO_2_/HCO_3_^−^ assimilation steps, the EMC pathway yields a 33% higher carbon retention per mole of assimilated acetyl-CoA. By incorporating these carboxylation sequences directly into the central carbon metabolism, the EMC pathway inherently functions as a metabolic electron sink, capitalizing on CO_2_ and HCO_3_^−^ reassimilation to accommodate intracellular electron surpluses while simultaneously maximizing carbon retention. More broadly, this architectural logic explains how non-photosynthetic lineages maximize carbon conservation and avoid redox bottlenecks during the assimilation of highly reduced, electron-dense substrates, such as methane and methanol.

Ultimately, the physiological utility of these intracellular architectures—whether operating as polymer buffers or carboxylating electron sinks—is tested by the dynamic fluctuations of the external environment. Systems featuring cyclic anaerobic and aerobic phases represent prime ecological arenas where competing microbial guilds must continuously deploy internal metabolic buffers to navigate shifting redox pressures and variable substrate availability. A classic exemplar of such dynamic selection is the enhanced biological phosphorus removal (EBPR) process, where two primary heterotrophic lineages— polyphosphate-accumulating organisms (PAOs) and glycogen-accumulating organisms (GAOs)—coexist in direct competition for shared resources within the same ecological niche. While real-world anoxic/anaerobic fermentation environments naturally co-generate formate and hydrogen (H_2_)^23^ alongside volatile fatty acids (VFAs), laboratory-scale EBPR investigations have overwhelmingly relied on simplified synthetic feeds containing acetate or propionate as sole carbon sources^24,25,26,27^. This conventional emphasis on VFA-exclusive feeding leaves a fundamental gap in our understanding of how concurrent energy and electron inputs from formate and H_2_ influence intracellular redox management, alter bioenergetic parities, and ultimately govern PAO–GAO competitive dynamics.

Parallel to this uncharacterized co-substrate physiology is a well-established kinetic paradigm dictating that competitive outcomes in EBPR systems are governed predominantly by instantaneous resource flux, driven by specific kinetic velocity and absolute biomass abundance^28–31^. While these rate-centric frameworks provide elegant descriptions of short-term anoxic substrate sequestration, standard steady-state assumptions often overlook how less obvious stoichiometric advantages derived from minor co-substrates translate into fractional yield gains. When biomass retention time is systematically decoupled from rapid hydraulic cycles, this marginal yield advantage can scale non-linearly over time, ultimately reshaping long-term community structure.

In pioneering long-term microbial evolution experiments, Lenski and colleagues^32^ formalized how even a fractionally higher growth yield or shorter lag phase—a minuscule fitness differential (*W*)—compounds exponentially over time. Over hundreds of serial transfer cycles, an advantage as small as 1% (*W* = 1.01) can result in the complete replacement of ancestral strain from a population. Such compounding effects are highly relevant in engineered processes where the solids retention time (SRT) is largely decoupled from the much faster, hours-long hydraulic retention time (HRT). This structural decoupling creates a venue for ‘metabolic compound interest’: a minute single-cycle stoichiometric advantage can snowball non-linearly over successive cycles, ultimately dictating which sub-population phenotypes persist.

In this study, using a well-controlled EBPR enrichment culture, we demonstrate how parallel strategies of inorganic resource management integrated with intracellular polymer dynamics establish a delicate bioenergetic parity between competing heterotrophic groups, and how minor fermentative co-substrates break this parity. Aerobically, GAOs deploy the EMC pathway for heterotrophic CO_2_/HCO_3_^−^ fixation during the mobilization of PHA reserves, utilizing inorganic carbon as a redox safety valve to absorb excess reducing equivalents and double carbon conservation efficiency in terms of biomass yield. Embedded within a highly synchronized metabolic topology, this EMC-driven stoichiometry balances the bioenergetic footing between GAOs and PAOs, shifting their competition into a classic rate-yield trade-off ^33–36^. This mechanistic revision fundamentally challenges a 40-year-old basic assumption regarding GAO bioenergetic inferiority in wastewater microbial ecology. Crucially, we show that minor formate co-feeding provides a minuscule per-cycle energetic advantage exclusively to GAOs. When operated under systematically decoupled biomass and hydraulic retention times, this marginal stoichiometric gain acts as “metabolic compound interest”, snowballing over three SRTs to drive a complete and irreversible population shift toward *Ca.* Competibacteraceae GAO dominance.

While demonstrated here within an engineered bioprocess, these findings reveal broader principles governing microbial niche differentiation and ecosystem stability across natural biofilms^36^ and attached-growth environments^37,38^. In ecosystems where biomass retention time is decoupled from fluid residence times, evolutionary selection does not exclusively favor sheer substrate uptake velocity (*q*_max_); instead, physical retention of biomass during rapid fluid throughput introduces generational compounding, creating a selective regime that favors long-term stoichiometric yield (*Y*) and elevated carbon-use efficiency^33,39^. Furthermore, our results show that functional niche differentiation among heterotrophs depends not only on competition for primary organic substrates, but also on the strategic management of inorganic resources (e.g., phosphate, CO_2_/HCO_3_^−^) and secondary co-substrates (e.g., formate, H_2_). Ultimately, this work offers a fresh mechanistic perspective on the long-standing puzzle of macro-scale EBPR performance instabilities^40^, while showcasing how minor environmental inputs break metabolic symmetries to shape microbial community assembly.

## 2. Materials and Methods

### 2.1 Bioreactor operations and configuration

A laboratory scale sequencing batch reactor (SBR) with a working volume of 1.5 L was operated in this study. The reactor was equipped with dedicated temperature, pH, and dissolved oxygen (DO) sensors, and was connected to an online gas chromatography-mass spectrometry (GC-MS) system for continuous off-gas analysis (H_2_, CO_2_, O_2_, N_2,_ Ar, and CH_4_). The reactor temperature was maintained at 19 ± 1 °C via ambient laboratory temperature control. Reactor pH was automatically regulated at 7.0 ± 0.2 via the dropwise addition of 0.5 M HCl and 0.5 M NaOH solutions.

Anaerobic/anoxic and aerobic conditions were maintained during their respective phases via the automated supply of either nitrogen gas or compressed air at a flow rate of 500 ml /min. Online off-gas analysis revealed that the exhaust gas O_2_ content fluctuated between 3.9% (during peak aerobic respiration) and 20.6% (toward the end of the famine phase). Liquid-phase DO was monitored via an online probe calibrated to air saturation (100%); DO levels dropped transiently to approximately 20% during peak initial PHA respiration before returning to near-saturation (∼100%) toward the end of the aerobic phase concurrent with PHA depletion.

The SBR was inoculated with aerobic granular sludge collected from the Utrecht wastewater treatment plant (Utrecht, The Netherlands). The reactor was operated continuously for over 9 months with an overall HRT of 9.6 h based on a 4.8 h cycle configuration. Each cycle consisted of an 11 min anaerobic filling period (during which 0.75 L of synthetic wastewater was introduced), a 120 min anaerobic phase, a 140 min aerobic phase; a 2 min sludge discharge phase, a 10 min settling phase, and a 5 min effluent withdrawal phase (0.75 L effluent discharged). To maintain a constant SRT of approximately 10 d, a fixed volume of approximately 30 mL of the mixed liquor suspended solids (MLSS) was purged during the sludge discharge phase prior to settling.

The influent medium was formulated to provide specific target concentrations of macronutrients and nitrification inhibitors in the synthetic wastewater: 922 mg/L CH_3_COONa•3H_2_O (434 mg COD/L), 82.7 mg/L NH_4_Cl (21.6 mg NH_4_-N/L), 136 mg/L NaH_2_PO_4_•2H_2_O (27 mgPO_4_-P/L), 86 mg/L MgSO_4_•7H_2_O, 22 mg/L CaCl_2_•2H_2_O, 26 mg KCl, 2 mg/L yeast extract, and 3.1 mg/L allylthiourea to achieve complete inhibition of nitrification. Additionally, the medium was supplemented with 300 µL/L of trace element solution prepared according to Smolders et al.^29^.

### 2.2 Experimental timeline and analytical sampling strategy

The experimental timeline was divided into two distinct operational phases. During Phase 1 (the baseline period, FA00), acetate was supplied as the sole carbon source (9 mM) to enrich a representative EBPR consortium. In Phase 2 (the co-feeding period), formate was introduced as a fermentative co-substrate, where the influent formate-to-acetate molar ratio was incrementally stepped up from 0.1:1 (transient phase FA01) to 0.2:1 (FA02).

Mixed liquor suspended solids (MLSS) and mixed liquor volatile suspended solids (MLVSS) were quantified weekly at the end of the anaerobic and aerobic phases to monitor biomass dynamics. These measurements were complemented by weekly tracking of cyclic concentration profiles for soluble orthophosphate (PO_4_^3-^), acetate and formate. A pseudo-steady state was defined and verified for each operational phase once macro-stoichiometric profiles and biomass concentrations remained stable for a minimum of three SRTs (≈30 days). Comprehensive full-cycle profiles of intracellular storage polymers—including glycogen and PHAs (specifically polyhydroxybutyrate [PHB] and polyhydroxyvalerate [PHV])—were executed once the system attained pseudo-steady state in each experimental phase.

Biomass samples were harvested during each full-cycle analysis and preserved for fluorescence *in situ* hybridization (FISH), metagenomic, and metaproteomic analyses. Oligonucleotide probes, staining conditions, and FISH visualization protocols are detailed in the Supporting Information (Section S1). To reconstruct the metabolic potential and enzyme expression profiles of the enriched consortia, metagenomic binning and metaproteomic quantification were performed. The complete computational and analytical workflow— including DNA/protein extraction protocols, sequencing strategies, *de novo* assembly, genome binning, protein identification, and bioinformatic pipeline parameters—is detailed in the Supporting Information (Section S2).

### 2.3 Physico-chemical analyses, stoichiometry and kinetic evaluation

All liquid samples collected for the quantification of soluble analytes were immediately clarified through 0.22 µm pore-size membrane filters. Intracellular PHA (PHB and PHV) and glycogen were extracted and quantified using established protocols described previously^29,41^. Soluble orthophosphate (PO_4_^3-^-P), and ammonium (NH_4_^+^-N) concentrations were determined colorimetrically using an automated discrete analyzer. Acetate and formate concentrations were resolved and quantified via an ultra-high performance liquid chromatography (UHPLC) system (Vanquish, Thermo Fisher Scientific, Breda, The Netherlands).

Key stoichiometric parameters and kinetic rates (Table 1) were calculated based on the observed net metabolite conversions across the anaerobic and aerobic phases. Specifically, these parameters included the anaerobic phosphate release-to-acetate uptake ratio (P-mol/C-mol), PHA stored per acetate consumed (C-mol/C-mol), glycogen degraded per acetate consumed (C-mol/C-mol), and the relative fractions of PHB and PHV within the total PHA pool^25^. The observed active biomass yield was estimated by subtracting the measured PHA and glycogen contents from routine MLVSS data, with the residual active biomass (*X*_active_) assigned an empirical elemental composition of C:H_1.77_:O_0.49_:N_0.24_^42^.

**Table 1.**
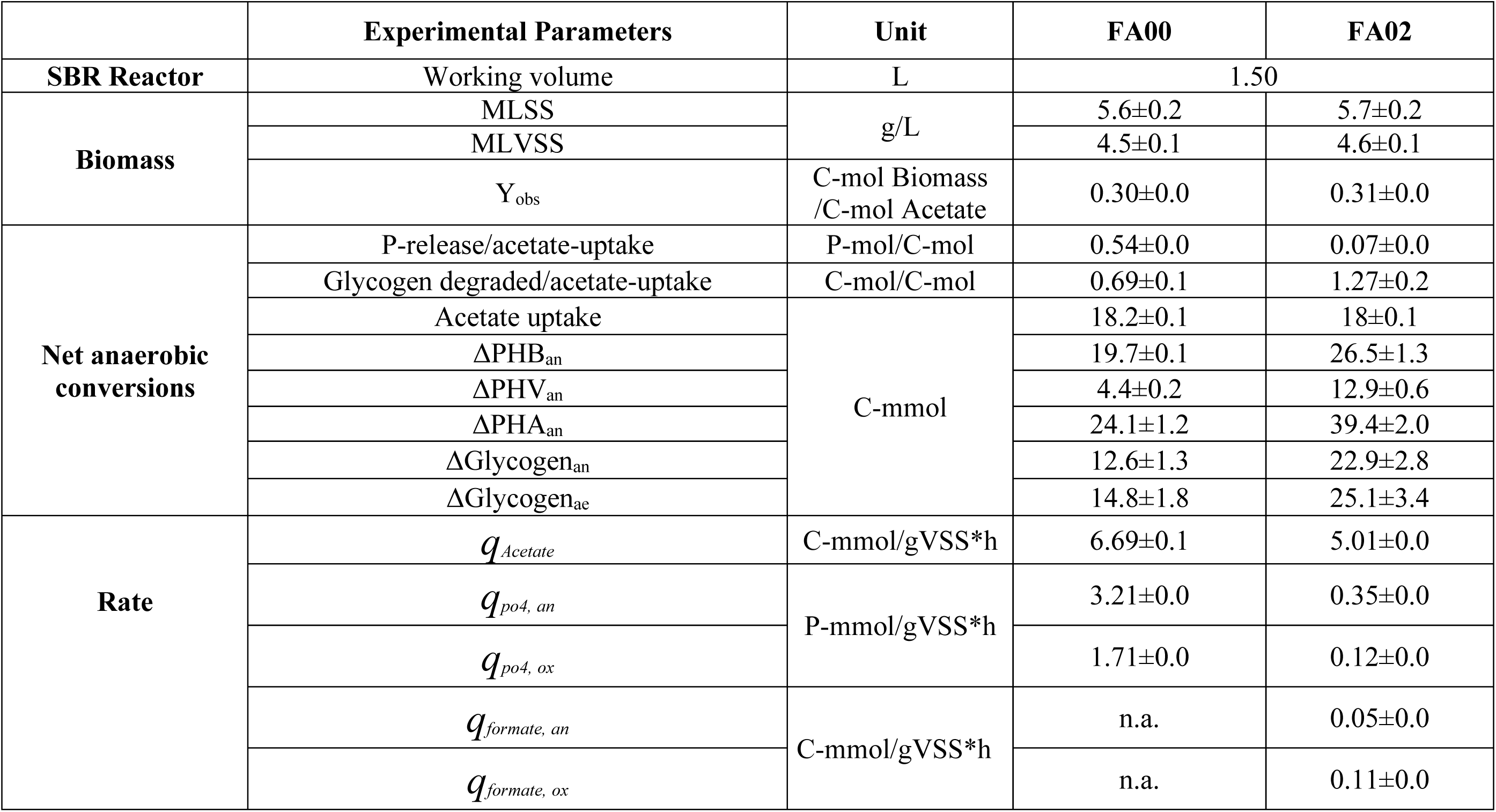
Stoichiometry and biomass specific rates observed under pseudo-steady-state conditions in EBPR enrichment in an anaerobic-aerobic SBR process, with acetate as carbon source (Experimental Phase 1 - FA00), and with formate as co-substrate (Experimental Phase 2 - FA02).

|  | Experimental Parameters | Unit | FA00 | FA02 |
| --- | --- | --- | --- | --- |
| **SBR Reactor** | Working volume | L | 1.50 |  |
| **Biomass** | MLSS | g/L | 5.6±0.2 | 5.7±0.2 |
|  | MLVSS |  | 4.5±0.1 | 4.6±0.1 |
| | $Y_{obs}$ | C-mol Biomass /C-mol Acetate | 0.30±0.0 | 0.31±0.0 |
| **Net anaerobic conversions** | P-release/acetate-uptake | P-mol/C-mol | 0.54±0.0 | 0.07±0.0 |
|  | Glycogen degraded/acetate-uptake | C-mol/C-mol | 0.69±0.1 | 1.27±0.2 |
|  | Acetate uptake | C-mmol | 18.2±0.1 | 18±0.1 |
| | $\Delta PHB_{an}$ | | 19.7±0.1 | 26.5±1.3 |
| | $\Delta PHV_{an}$ | | 4.4±0.2 | 12.9±0.6 |
| | $\Delta PHA_{an}$ | | 24.1±1.2 | 39.4±2.0 |
| | $\Delta Glycogen_{an}$ | | 12.6±1.3 | 22.9±2.8 |
| | $\Delta Glycogen_{ae}$ | | 14.8±1.8 | 25.1±3.4 |
| **Rate** | $q_{Acetate}$ | C-mmol/gVSS*h | 6.69±0.1 | 5.01±0.0 |
| | $q_{po4, an}$ | P-mmol/gVSS*h | 3.21±0.0 | 0.35±0.0 |
| | $q_{po4, ox}$ | | 1.71±0.0 | 0.12±0.0 |
| | $q_{formate, an}$ | C-mmol/gVSS*h | n.a. | 0.05±0.0 |
| | $q_{formate, ox}$ | | n.a. | 0.11±0.0 |

## 3. Results

### 3.1 Formate-driven shift from *Ca.* Dechloromonas PAO-dominated community to *Ca*. Competibacter GAO-dominated community and deterioration of EBPR performance

To investigate the physiological impact of the fermentation byproduct, specifically formate, on EBPR performance, a baseline EBPR consortium was first established under standard acetate-only feeding until reaching pseudo-steady state (FA00). We then introduced a minor amount of formate as a secondary co-substrate to evaluate its impact on community succession, macro-stoichiometry, and phosphorus removal capacity. Specifically, following a brief transition period (FA01) with low-dose formate co-feeding (formate:acetate = 0.1:1, mol:mol), we operated the reactor under a 0.2:1 (mol:mol) formate:acetate regime (FA02) until reaching a new pseudo-steady state.

During the formate-free baseline phase (FA00), with acetate as the sole carbon source, the system exhibited robust, classical EBPR performance driven by a co-dominant assembly of *Ca.* Dechloromonas PAO and *Ca.* Competibacteraceae GAO (Table 1, Figure 1, Figure 2, Figure S2). Under these baseline conditions, the anaerobic phosphate release-to-acetate uptake ratio averaged 0.54±0.2 P-mol/C-mol, with a specific anaerobic phosphate release rate (*q*_PO4,an_) of 3.21±0.4 P-mmol gVSS⁻¹ h⁻¹.

**Figure 1.**
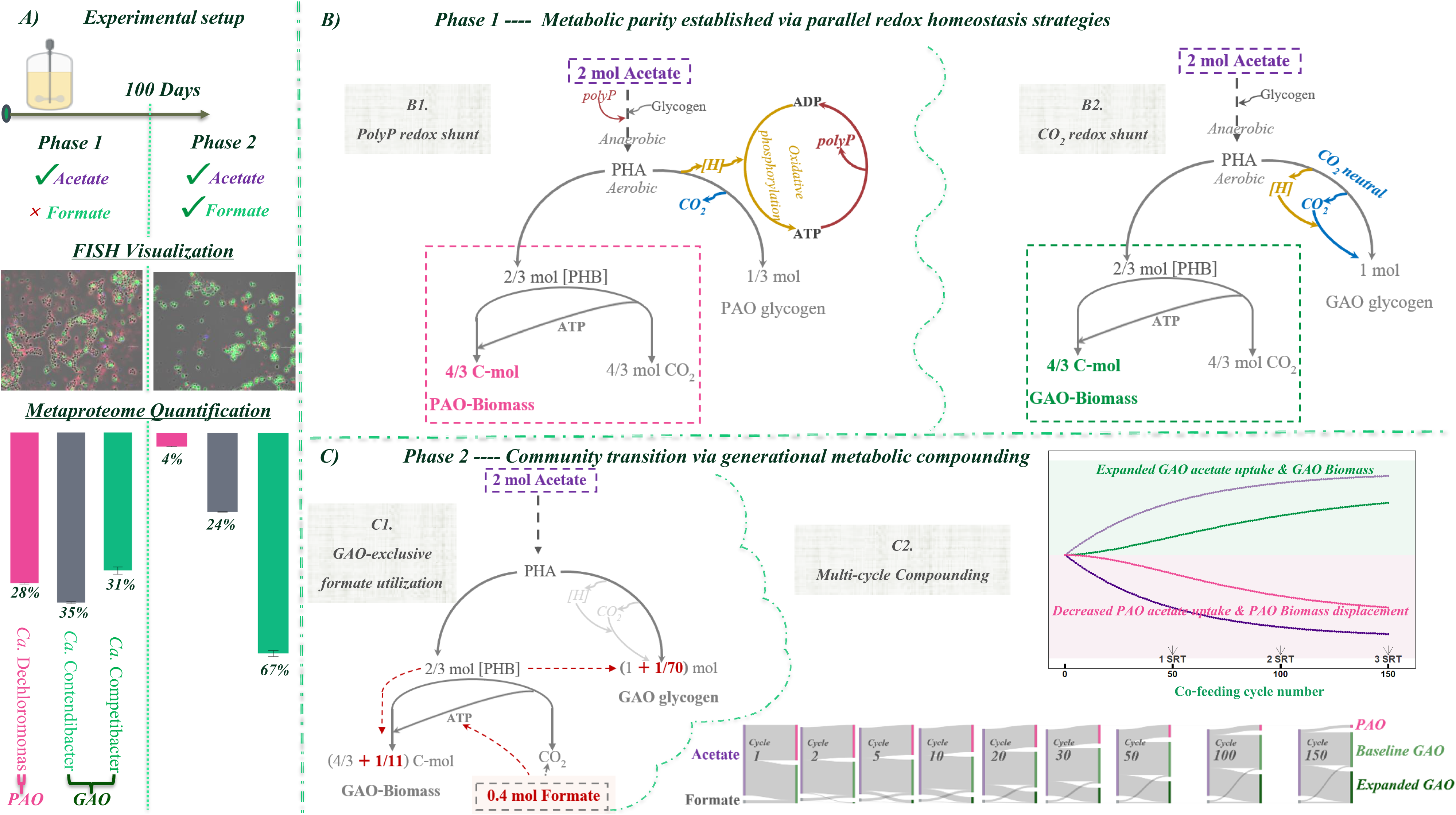
Conceptual overview of baseline PAO–GAO metabolic parity and formate-induced competitive displacement. **(A) Experimental layout and community profiling.** Overview of reactor operation across Phase 1 (acetate baseline) and Phase 2 (acetate + formate co-feeding), validated by FISH microscopy and metaproteomic quantification (see Figure 2 and Figure S2). **(B) Phase 1 (FA00): Baseline metabolic parity.** Stoichiometric accounting demonstrates functional metabolic parity under baseline conditions: starting from 2 mol of anaerobic acetate uptake, both polyphosphate-mediated (*B1*, *Ca.* Dechloromonas PAO) and CO_2_-mediated (*B2*, *Ca.* Competibacteraceae GAO) strategies allocate an identical 2/3 mol of PHB toward aerobic biomass synthesis, yielding equal growth yields (see Figure 6B and Figure 6C for quantitative carbon/energy flux maps). **(C) Phase 2: Community restructuring via generational metabolic compounding.** GAO-exclusive formate utilization (*C1*) yields boosts aerobic GAO biomass synthesis (by + 1/11 C-mol) and GAO glycogen storage (by + 1/70 mol). This asymmetric energetic advantage initiates a multi-cycle compounding feedback loop (*C2*), driving substrate monopoly and exponential PAO displacement over 150 cycles / 3 SRTs (see Figure 7 for the complete quantitative simulation).

**Figure 2.**
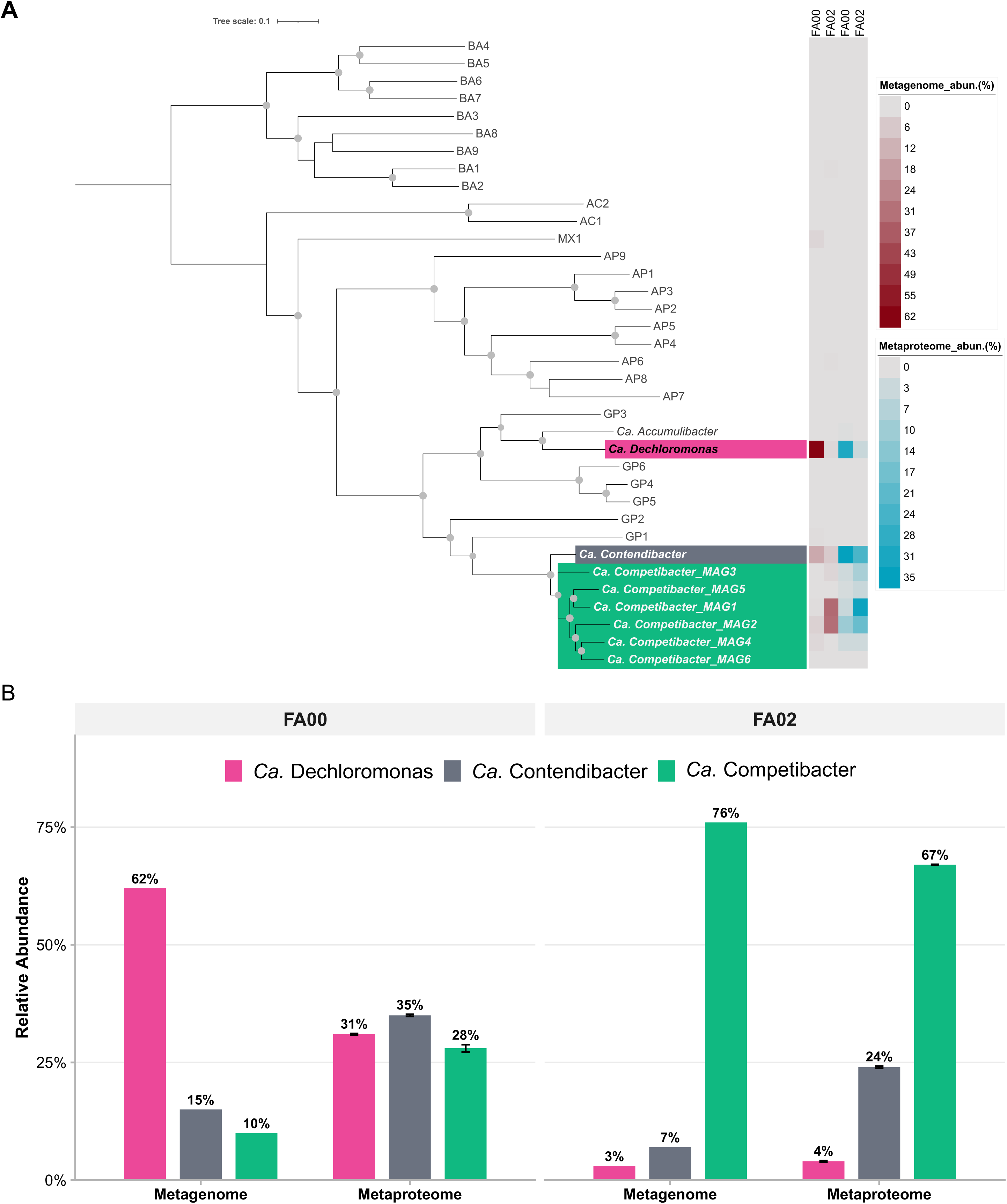
Phylogenomic resolution and multi-omics profiling of recovered metagenome-assembled genomes. **(A) Maximum-likelihood phylogenomic tree.** Inferred from a concatenated alignment of 120 ubiquitous single-copy bacterial marker proteins across 35 recovered draft metagenome-assembled genomes (MAGs). Mapped heatmaps display the relative abundance of each genome at the metagenomic DNA level (red) and metaproteomic expression level (blue) across experimental phases FA00 and FA02. Scale bar represents 0.1 substitutions per amino acid site. Detailed genomic metrics for all 35 MAGs are provided in Supplementary Appendix File 1**. (B) Comparative metagenomic vs. metaproteomic quantification.** Relative abundance of the dominant *Ca.* Dechloromonas (PAO), *Ca.* Contendibacter (GAO), and *Ca.* Competibacter (GAO) populations. Discrepancies between DNA- and protein-based quantification highlight a widespread but underappreciated bias in community profiling: metagenomic sequencing overrepresents populations with smaller cell size, whereas metaproteomics better reflects true biomass volume. Specifically, in this study, a metagenome-based PAO dominance at >60% in FA00 translates to a combined GAO dominance at the metaproteome level, where *Ca.* Dechloromonas PAO (∼31%) and the two GAO genera (∼35% *Contendibacter* and ∼28% *Competibacter*) each represent approximately one-third of the total detected peptide spectra.

Transitioning to the 0.2:1 (mol:mol) formate-to-acetate co-feeding regime (FA02) triggered a rapid community restructuring. Within just three SRTs, competitive exclusion led to a *Ca.* Competibacter–dominated culture, coinciding with a total collapse of EBPR activity (Figure 1, Table 1). Stoichiometrically, the system shifted from a polyphosphate-dependent PAO phenotype during baseline (FA00; 0.54±0.2 P-mol/C-mol) to a classical glycogen-fueled GAO phenotype in FA02, where phosphate release virtually ceased (0.07±0.0 P-mol/C-mol). Concurrently, glycogen flux nearly doubled to compensate for the lost polyphosphate energy pool, directly fuelling increased synthesis of total PHA and a higher portion of the PHV fraction (Table 1, Figure S1). This swift transition demonstrates that minor formate co-feeding completely disengages polyphosphate reliance in favor of a glycogen-driven GAO metabolism.

To understand why such a minor co-substrate addition could trigger a complete community reversal, we first examined how *Ca.* Competibacteraceae GAOs and *Ca.* Dechloromonas PAOs handle formate at the metabolic level.

### 3.2 Divergent formate-metabolizing potential and H_2_-handling strategies among PAO and GAO genotypes

With the absence of both the catabolic formate dehydrogenases (*fdh*) and the anabolic formyltetrahydrofolate synthetase (*fhs*), genome annotation and metaproteomic profiling revealed that formate utilization is mediated exclusively via the membrane-bound formate hydrogenlyase (*FHL*-2) complex^43,44^ in two members of the *Ca.* Competibacteraceae, converting formate to H_2_, CO_2_, and ATP (Table S1). In contrast, *Ca.* Dechloromonas PAO remained largely inactive toward formate. Downstream H_2_ utilization was linked to two distinct hydrogenases: a cytoplasmic, NAD-reducing hydrogenase (*Hox*)^45^ mediating reversible electron transfer between H_2_ and the NAD^+^/NADH pool, and a membrane-bound respiratory NiFeSe hydrogenase (*Hyd*-2; Table S1)^46,47^. Robust metaproteomic expression of *Hyd*-2, combined with the rapid drawdown of accumulated H_2_ to undetectable levels upon aeration (Figure 3), strongly points to aerobic H_2_ oxidation via quinone-coupled electron transport as a primary mechanism for respiratory ATP generation.

**Figure 3.**
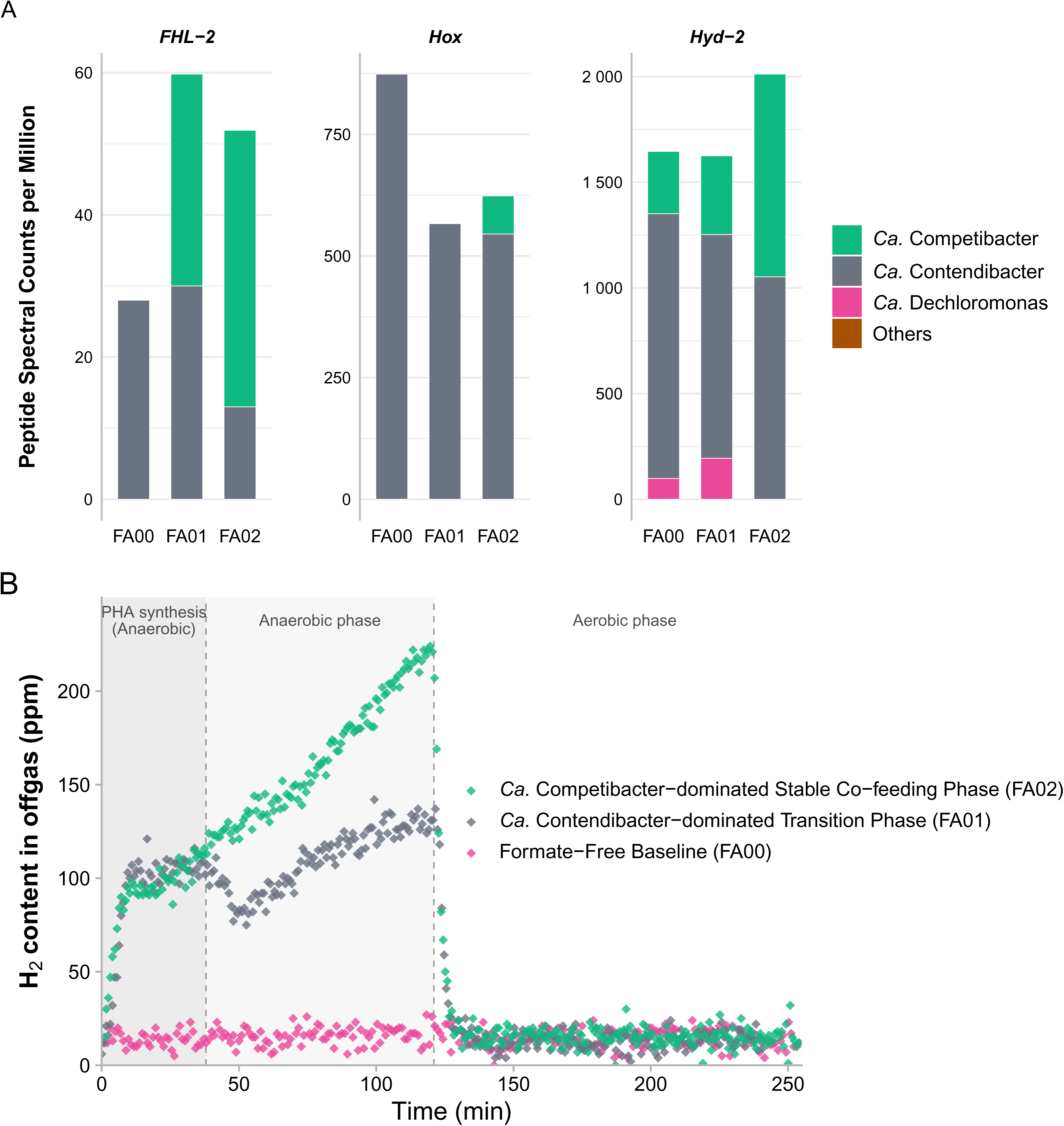
Metaproteomic profiles of core enzyme modules involved in formate & H_2_ metabolism, and real-time off-gas H_2_ dynamics across operational phases. **(A) Metaproteomic expression profiles of the core enzyme modules** expressed as peptide spectral counts per million matched spectra across three distinct experimental phases: the formate-free baseline (FA00), the early co-feeding transition phase dominated by *Ca.* Contendibacter GAO (FA01), and the late co-feeding phase dominated by *Ca.* Competibacter GAO (FA02). *FHL-2*: Formate-hydrogen lyase complex driving formate dismutation to H_2_; *Hox*: Cytoplasmic NiFeSe-hydrogenase mediating electron transfer from H_2_ to NADH; *Hyd-2*: Membrane-bound respiratory NiFeSe-hydrogenase coupled to aerobic respiration. Stacked bars designate taxonomic assignment down to the genus level (*Ca.* Contendibacter, *Ca.* Competibacter, and *Ca.* Dechloromonas). The negligible detection of these enzyme modules in flanking community members (“Others”) confirms that Ca. Competibacteraceae GAOs dominant the formate and H_2_ metabolism (see Table S2 for a detailed biochemical breakdown of each enzyme module). **(B) Real-time online off-gas H_2_ dynamics during individual SBR cycles.** Headspace H_2_ concentration profiles across the formate-free baseline (FA00, pink), the *Ca.* Contendibacter-dominated early transition phase (FA01, grey), and the *Ca.* Competibacter-dominated stable co-feeding phase (FA02, teal). The pronounced accumulation of headspace H_2_ during the first 120 minutes corresponds to the anaerobic reaction period, directly reflecting *in vivo* formate-hydrogen lyase (*FHL-2*) activity. The abrupt collapse at 120 min marks the onset of aeration, where accumulated H_2_ is rapidly depleted via respiratory oxidation.

**Figure 4.**
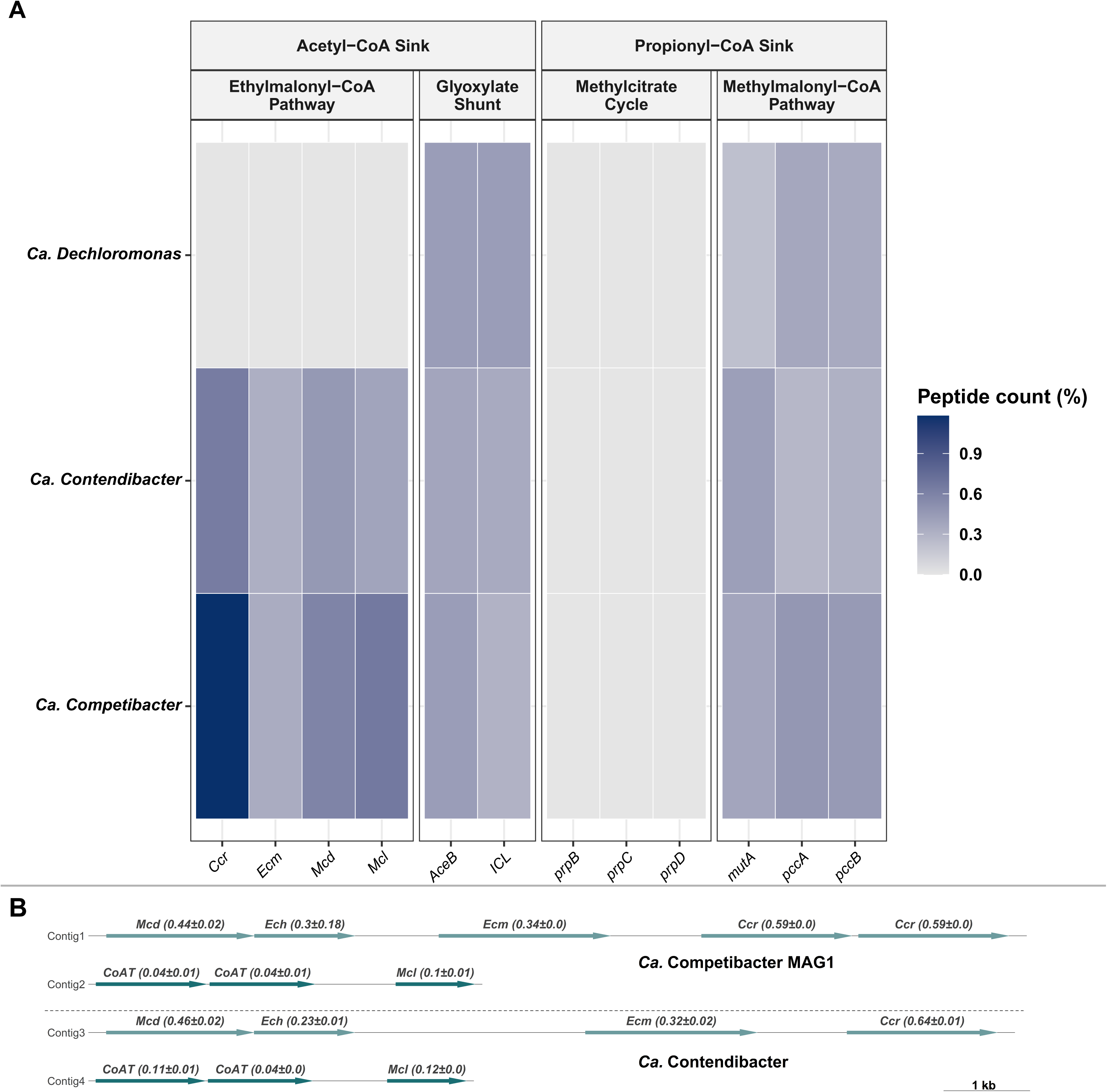
Comparative metaproteomic profiling of central carbon metabolism sinks and genomic organization of the ethylmalonyl-CoA (EMC) gene cluster. **(A) Metaproteomic heatmap of central carbon pathways.** Relative peptide abundance (%) of key enzymes involved in acetyl-CoA and propionyl-CoA assimilation across *Ca.* Dechloromonas, *Ca.* Contendibacter, and *Ca.* Competibacter MAG1. Both GAO clades exclusively express the Ethylmalonyl-CoA Pathway (*Ccr*, crotonyl-CoA reductase; *Ecm*, ethylmalonyl-CoA mutase; *Mcd*, methylsuccinyl-CoA dehydrogenase; *Mcl*, malyl-CoA lyase) for dark CO_2_fixation and redox homeostasis, whereas *Ca.* Dechloromonas exclusively utilizes the Glyoxylate Shunt (*AceB*, malate synthase; *ICL*, isocitrate lyase). **(B) Genomic synteny and active translation of the EMC gene cluster.** Conserved operon organization of EMC pathway genes across recovered metagenome-assembled genomes (MAGs) of *Ca.* Competibacter (Contigs 1–2) and *Ca.* Contendibacter (Contigs 3–4). Values above arrows denote relative peptide abundance (mean ± SD). Scale bar = 1 kb.

While both *Ca.* Contendibacter GAO and *Ca.* Competibacter GAO actively consumed formate, they displayed a clear divergence in anaerobic H_2_ management. During anaerobic PHA synthesis, *Ca.* Contendibacter recycled H_2_-derived electrons back into the intracellular electron pool via *Hox*, generating NADH that can serve as the required reducing equivalent for PHA biosynthesis; whereas *Ca.* Competibacter maintained minimal *Hox* expression (Figure 3). Real-time off-gas dynamics independently validated this divergence. During the *Ca.* Contendibacter– dominated phase (FA01), anaerobic H_2_ concentrations plateaued at ≈104 ppm (E’_H_^+^_/H_2 = -290 mV) during PHA synthesis, tightly poising the system near the physiological NAD^+^/NADH midpoint potential (-280 mV)^48^ and reflecting active intracellular H_2_ re-oxidation via *Hox* (Figure 3). Conversely, during the *Ca.* Competibacter–dominated phase (FA02), off-gas monitoring revealed continuous, linear H_2_ accumulation beyond 200 ppm. This continuous venting reflects unmitigated H_2_ evolution by *FHL-2* that exceeds the re-consumption capacity of the relatively low expressed *Hox* complex in Ca. Competibacter.

This metabolic divergence explains the shift in GAO microdiversity from *Ca.* Contendibacter-*Ca.* Competibacter co-dominance to *Ca.* Competibacter dominance. Because standard GAO metabolism already relies on the reductive branch of the TCA cycle to sink the excess NADH generated during glycolysis for PHA synthesis (Figure S4), additional NADH generation via *Hox*-mediated H_2_ re-oxidation imposes an extra redox penalty on *Ca.* Contendibacter, requiring elevated glycogen expenditure for electron balance. In contrast, *Ca.* Competibacter captures metabolic energy from formate via *FHL-2* while safely venting excess reducing equivalents as H_2_ gas during the anaerobic phase, thereby avoiding redox stress and outgrow *Ca.* Contendibacter under sustained formate co-feeding.

While this divergence explains how *Ca*. Competibacteraceae GAOs handle formate once available, it does not explain why GAOs were positioned to outcompete PAOs, given the recognized kinetic competitiveness of PAOs under standard VFA feeding^9^. Addressing this requires re-examining how the two guilds achieved competitive parity in the first place.

### 3.3 Metabolic divergence in central carbon metabolism: the glyoxylate shunt versus the ethylmalonyl-CoA pathway

During aerobic PHA mobilization, both PAOs and GAOs face an intracellular redox challenge arising from the high-flux oxidation of reduced PHA reserves toward glycogen. In canonical PAOs, excess reducing equivalents generated during PHA-to-glycogen routing are dissipated, in part, via energetic coupling to polyphosphate synthesis (Figure 1). Conversely, GAOs lack polyphosphate storage while simultaneously requiring a threefold higher carbon flux from PHA to replenish glycogen. This massive routing causes a threefold higher electron overflow and substantial CO_2_ loss from localized decarboxylation (Figure 6a), creating a severe stoichiometric mismatch: an abundance of electrons paired with a shortage of carbon skeletons for biomass synthesis. Rather than relying on the glyoxylate shunt like PAOs, GAOs circumvent this constraint through heterotrophic, reductive CO_2_/HCO_3_^−^ assimilation embedded within the EMC pathway (Figure 1, Figure 6b, Figure 5).

**Figure 5.**
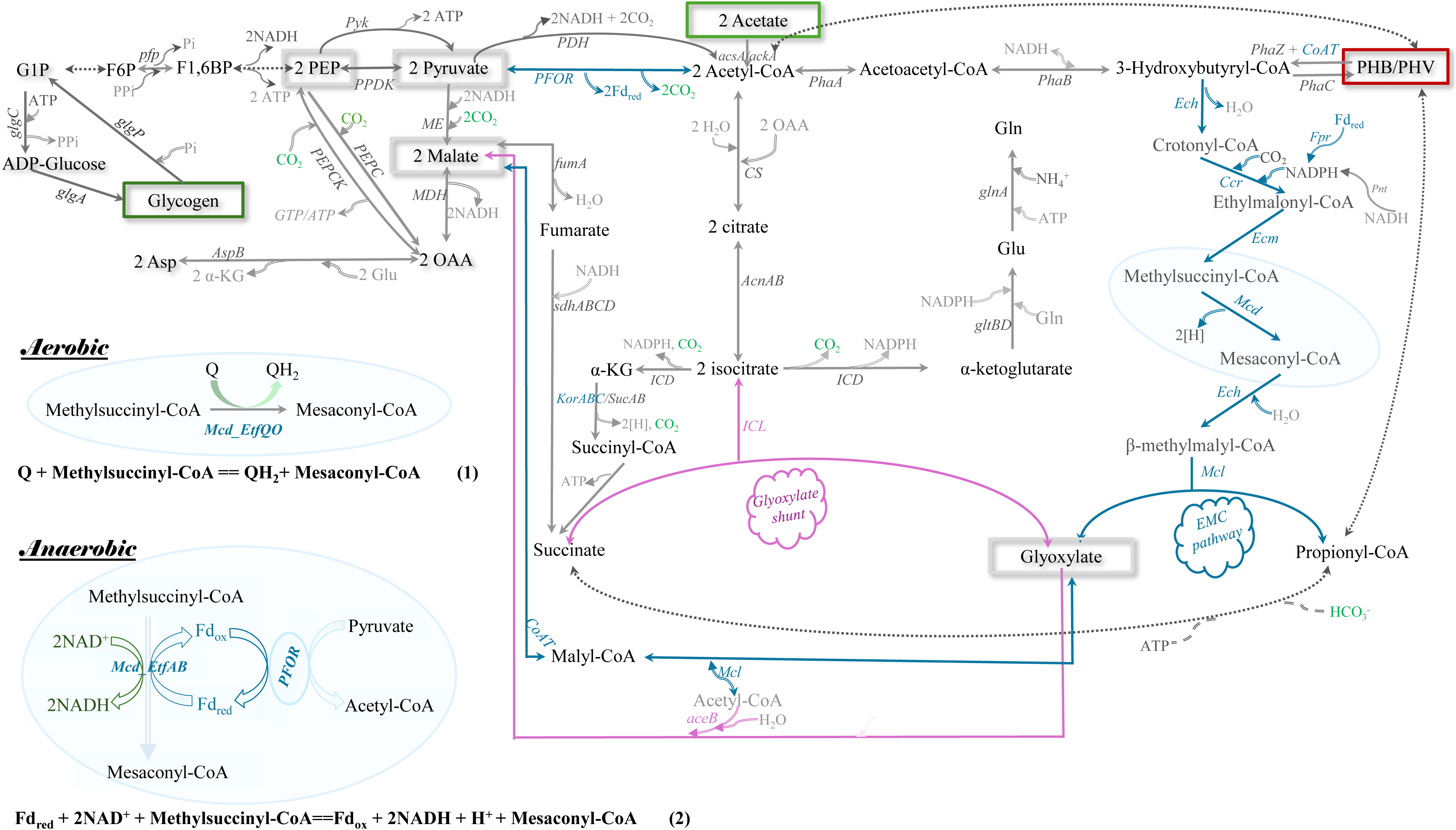
Comprehensive metabolic network topology of central carbon metabolism in *Ca.* Competibacteraceae. Schematic reconstruction highlights the metabolic wiring driving glycogen and polyhydroxyalkanoate (PHA) interconversions, carbon routing through the Glyoxylate Shunt (pink lines) and the Ethylmalonyl-CoA (EMC) pathway (blue lines). Key enzymatic sub-modules, including pyruvate:ferredoxin oxidoreductase (*PFOR*) and ferredoxin-NADP^+^ reductase (*Fpr*), are highlighted in blue to denote their roles in generating reduced ferredoxin (Fd_red_) and NADPH. Left-hand callouts: Detailed biochemical mechanisms for the aerobic and hypothetical anaerobic oxidation of methylsuccinyl-CoA to mesaconyl-CoA. Under aerobic conditions (Eq. 1), the *Mcd–EtfQO* complex mediates direct electron transfer from cytosolic methylsuccinyl-CoA to the membrane-bound quinone pool. Under anaerobic conditions (Eq. 2), the *Mcd–EtfAB* complex is proposed to catalyse flavin-based electron confurcation (FBEB family): coupling the thermodynamically uphill electron transfer from methylsuccinyl-CoA to NAD^+^ with the thermodynamically downhill drive from Fd_red_ to NAD^+^. Dashed lines represent multi-step transformations. Complete stoichiometric matrices are detailed in Supplementary Appendix File 2. Reduced ferredoxin is treated as a two-electron carrier in this study.

**Figure 6.**
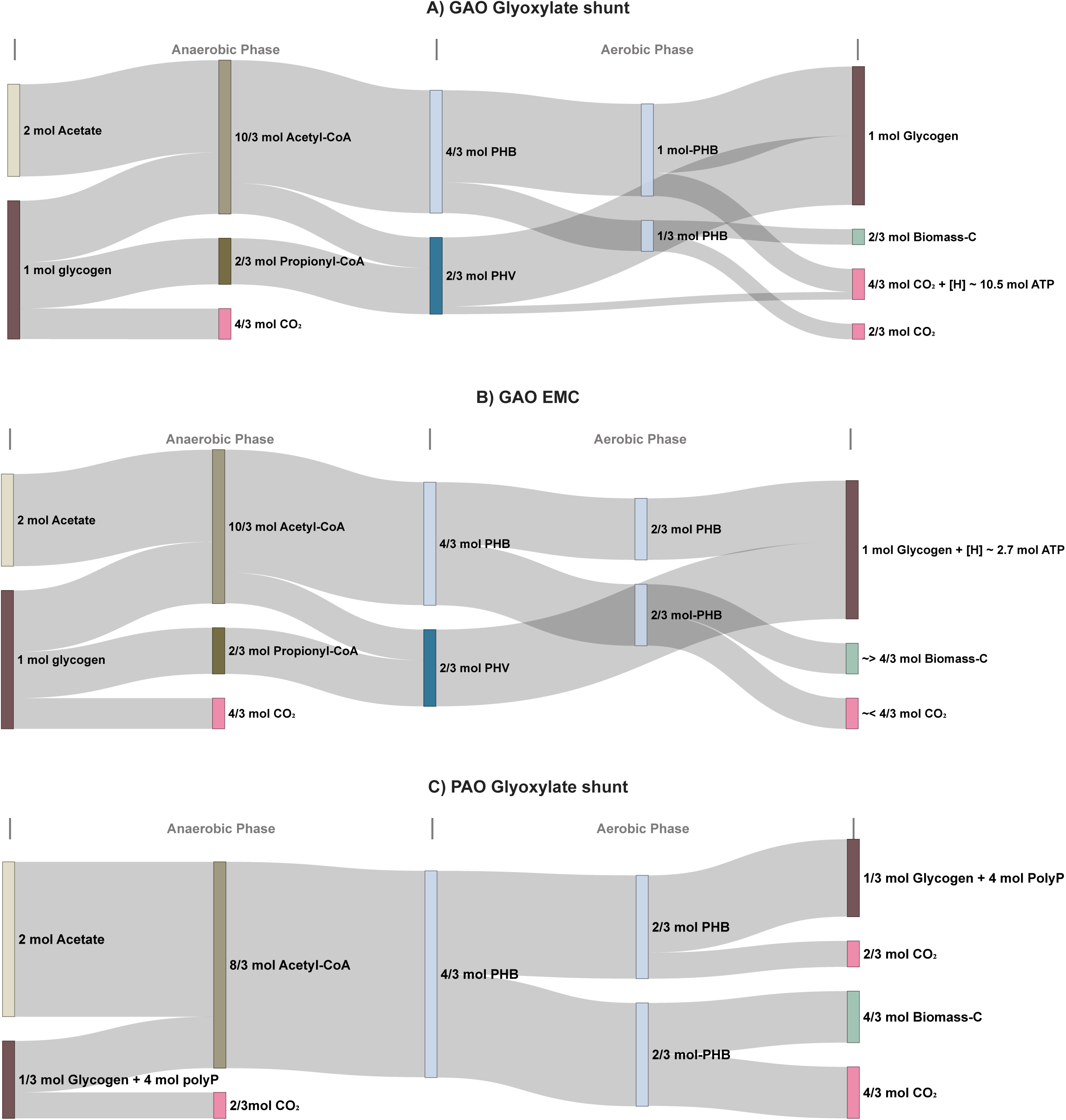
Quantitative stoichiometric flux maps contrasting carbon conservation efficiency in PAO and GAO phenotypes. Sankey diagrams track quantitative carbon routing and network stoichiometry throughout a complete anaerobic–aerobic cycle across three distinct metabolic modes: **(A)** GAOs utilizing a standalone Glyoxylate Shunt network (yielding a theoretical maximum of 0.16 C-mol biomass/C-mol acetate), **(B)** GAOs utilizing an integrated EMC pathway (yielding 0.33 C-mol biomass/C-mol acetate) and **(C)** canonical PAO phenotype utilizing the Glyoxylate Shunt. Horizontal ribbons map the precise partitioning of initial substrates (acetate and glycogen) into intracellular PHAs, specifically PHB (as 3-hydroxybutyrate monomer equivalents) and PHV (as 3-hydroxyvalerate monomer equivalents), and their subsequent aerobic allocation into replenishment, biomass synthesis, and CO_2_. Ribbon widths reflect flux densities derived from stoichiometric matrix calculations (Supplementary Appendix File 2; corresponding anaerobic boundaries are detailed in Figure S3). Due to qualitative variations in biomass precursor requirements, metabolic trends of enhanced carbon conservation via the EMC pathway are indicated using ‘∼>’ and ‘∼<’ symbols in panel B).

Genome annotation and metaproteomic revealed that *Ca.* Competibacter GAO harbours tandem crotonyl-CoA carboxylase/reductase (*Ccr*) genes (Figure 4B), collectively commanding ≈1.2% of total detected spectra in *Ca.* Competibacter GAO proteome. While *Ca.* Contendibacter GAO displays similar proteomic investment in EMC machinery, *Ca.* Dechloromonas PAO relies exclusively on the glyoxylate shunt (Figure 4A), maximizing electron flow toward the respiratory chain to generate ATP for polyphosphate replenishment (Figure 1).

Stoichiometric flux analysis demonstrates that oxic EMC activation elevates the carbon conservation efficiency of the GAO phenotype (Figure 6b). Starting from 2 mol assimilated acetate, a conventional GAO model restricted to glyoxylate shunt suffers obligatory carbon loss as CO_2_, leaving only 1/3 mol of PHB for biomass synthesis and capping theoretical biomass yield at ≈ 0.16 C-mol/ C-mol acetate (Figure 6a). In contrast, routing carbon through the oxic EMC pathway renders glycogen replenishment CO_2_-neutral and preserves 2/3 mol PHB for biomass synthesis, doubling theoretical yield to ≈ 0.33 C-mol/C-mol acetate (Figure 6b).

This EMC adaptation establishes near-exact stoichiometric yield parity between GAOs and glyoxylate shunt-operating PAOs (Figure 6c), with both preserving 2/3 mol of PHB for biomass synthesis (≈ 0.33 C-mol biomass/C-mol Acetate). This theoretical convergence aligns with empirical biomass yield reported in enriched cultures^39,49^ and observed in this study (Table 1). While the multi-branched biochemistry of structural biomass synthesis is beyond the scope of this study, the intrinsic carbon-conserving capacity of the EMC pathway likely cascades into broader anabolic networks, securing a more efficient carbon-routing landscape than in glyoxylate shunt–dependent PAOs.

These findings reveal a fundamental evolutionary divergence during aerobic PHA mobilization: PAOs utilize an inorganic phosphate cycle as their primary metabolic buffer, whereas GAOs exploit inorganic carbon (CO_2_/HCO_3_^−^) as an oxic metabolic buffer to simultaneously secure structural carbon conservation and mitigate intracellular electron overflow.

While this aerobic strategy explains GAOs’ equivalent biomass yield during PHA mobilization, PAOs, with their Poly-P reserves, retain an anaerobic kinetic advantage via rapid substrate uptake^9,29,50,51^. For co-substrates like formate to tilt the balance, GAOs require an equally efficient anaerobic strategy to convert a minor bioenergetic edge into long-term dominance.

### 3.4 Integrated bioenergetic modules support high-efficiency resource utilization in *Ca*. Competibacteraceae GAOs

#### 3.4.1 The *PPDK–AcsA* module: high-affinity acetate activation at minimal energetic cost

In *Ca.* Competibacter and *Ca.* Contendibacter, co-expression of pyruvate-phosphate dikinase (*PPDK*) and acetyl-CoA synthetase (*AcsA*) enables high-affinity, low-energy acetate activation (Figure S3a). The glycogen-to-acetate stoichiometry (∼1: 2, mol: mol; Figure 5b) establishes a 1:1 molar ratio between glycogen-derived phosphoenolpyruvate (PEP) flux to pyruvate via *PPDK* and acetate activation via *AcsA*. Utilizing PEP to recycle AMP and inorganic pyrophosphate (PPi) while regenerating ATP for *AcsA* halves the net energy investment for carbon assimilation compared to standalone *AcsA*^52^ (Figure S3c). Notably, *Ca.* Dechloromonas PAO also relies on high-affinity *AcsA* over the lower-affinity *AckA–Pta* system (Figure S6). However, lacking PPDK and operating at a lower glycogen-to-acetate ratio (∼1: 6, mol: mol; Figure 6c), PAOs incur a higher energetic cost for equivalent acetate affinity, unless smooth energy coupling is achieved via ion-translocating pyrophosphatases.

#### 3.4.2 Ferredoxin-centered redox poising and energy conservation

The anaerobic bioenergetics of GAOs are further fortified by a coordinated, ferredoxin-centered metabolic module. Deploying pyruvate:ferredoxin oxidoreductase (*PFOR)* during glycogen breakdown utilizes low-potential ferredoxin ^1^ (Fd_red_) as the primary electron carrier^53^—a feature absent in *Ca.* Dechloromonas PAO.

Metaproteomics confirmed robust co-expression of PFOR, ferredoxin-NADP^+^ reductase (*Fpr*) and proton-translocating NAD(P)^+^ transhydrogenase (*PntAB*)^54,55^ in GAOs. Routing Fd_red_-derived electrons through *Fpr* and *PntAB* to NADH drives membrane-bound ion translocation, conserving an additional 0.5 mol ATP per mole of Fd_red_ oxidized. Furthermore, Fd_red_ may power an anaerobic variant of the EMC pathway (Reaction 2, Figure 5); this working hypothesis is fully expanded in the Supporting Information (Section S3).

#### 3.4.3 Interconnected central carbon network optimized for flexible and energetically economical polymer recycling

Reconstructing the *Ca.* Competibacteraceae metabolic network reveals high topological connectivity optimized for bidirectional carbon routing between PHA and glycogen (Figure 5). Exclusive expression of phosphoenolpyruvate carboxykinase *(PEPCK*) over *PEPC*, alongside *PPDK*, *PFOR*, and PPi-dependent phosphofructokinase (*Pfp*), provides a fully reversible glycolytic backbone (Figure S6).

Complementing this upper reversible glycolytic backbone, co-expression of the glyoxylate shunt and the EMC pathway (Figure 4A) establishes a bidirectional highway linking glycogen and PHA pools. Specifically, while isocitrate lyase (*ICL)* permits the reversible channelling between acetyl-CoA and glyoxylate, the terminal section of the EMC pathway enables the reversible interconversion of glyoxylate to malate, with malate serving as a vital precursor for glycogen synthesis. This architectural integration ensures that metabolic routing between the two primary intracellular polymers—glycogen and PHAs—is never bottlenecked by rigid, unidirectional pathways (Figure 5). Furthermore, duplicate family III CoA-transferases (*CoAT*) embedded in the EMC clusters (Figure 4B) mediate ATP-independent substrate-level CoA recycling. By pairing exergonic malyl-CoA cleavage with endergonic hydroxybutyrate activation, GAOs drive PHB-to-malate conversions with zero net ATP investment.

This structural configuration represents a major evolutionary departure from classical models of the EMC pathway. As two dominant mechanisms dedicated to assimilating exogenous C_1_ or C_2_ substrates, the glyoxylate shunt and the EMC pathway are typically mutually exclusive^22^. However, *Ca.* Competibacteraceae GAOs defy this evolutionary exclusion by co-expressing both pathways, enabling flexible interconversion of intracellular reserves under dynamic feast–famine conditions, so they can adapt quickly to the “feast-famine” lifestyle and dynamic environments.

Together, these aerobic and anaerobic adaptations establish that GAOs achieve baseline stoichiometric parity with PAOs. Next, we demonstrate how supplementary bioenergetics from formate co-feeding converts this baseline parity into rapid GAO dominance within three SRTs.

### 3.5 Stoichiometric basis for GAO dominance under formate co-feeding

Stoichiometric parameters derived from metabolic network reconstruction (Appendix File 2) informed a dynamic multi-cycle model simulating the competitive displacement of *Ca.* Dechloromonas PAO by *Ca.* Competibacter GAO within three SRTs under formate co-feeding.

In phase FA02, 31% (0.56 mmol) of co-fed formate was consumed anaerobically and 69% (1.24 mmol) was carried over and oxidized aerobically (Figure S5). Respiratory oxidation of the carried-over formate yields approximately 3.1 mmol ATP per aerobic cycle (Table S3, Appendix File 3), directly sparing 0.14 mmol PHB otherwise required for respiration. This spared PHB fraction is redirected toward anabolic metabolism (Figure 6b), expanding the glycogen pool by 0.08 mmol and GAO biomass-C by 0.14 mmol per cycle.

Because HRT (9.6 hours) is decoupled from SRT (10 days), this single-cycle stoichiometric advantage accumulates within the reactor, as expanded glycogen reserves and GAO-biomass are largely retained rather than washed out (only ∼ 2% wastage per cycle). Following this constrained mathematical framework, the simulated net glycogen expansion of 2.1 mmol over 3 SRTs (Figure 7, Appendix file 3) provides an excellent fit with our experimental measurements, specifically a 1.7 ± 0.5 mmol glycogen pool expansion from phase FA00 to FA02 (Table 1, Figure S1). Furthermore, the simulated formate-induced GAO biomass accrual of 3.0 g closely mirrors the PAO-biomass shrinkage of 3.0 g, elegantly accounting for the clear phenotypic transition from initial GAO-PAO parity to complete GAO dominance, characterized by the displacement of *Ca.* Dechloromonas PAO by *Ca.* Competibacter GAO (Figure 1; Figure 2; Figure S2). Section S4 of the Supporting Information provides a detailed description of the stoichiometric simulation.

**Figure 7.**
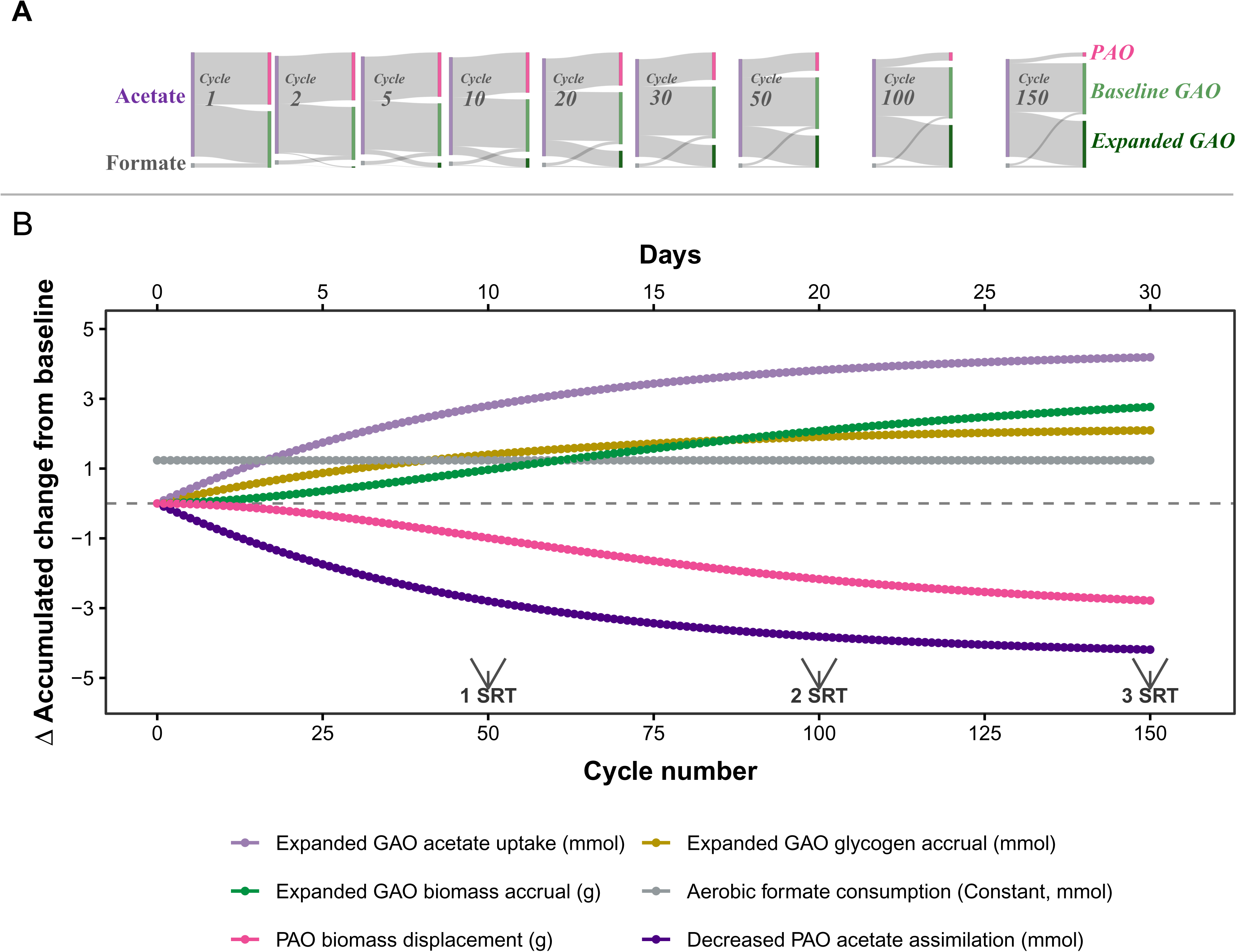
Generational metabolic compounding and progressive substrate monopoly in *Ca.* Competibacteraceae GAO over 150 operational cycles in 3 SRTs. **(A) Discrete multi-cycle flux allocations.** Stepwise Sankey diagrams track the dynamic reallocation of the anaerobic acetate feed across key cycles (Cycle 1 to 150), illustrating the shift from initial functional parity (50/50 PAO/GAO) to a progressive substrate monopoly driven by the expanded *Ca.* Competibacter GAO population under formate co-feeding. **(B) Continuous multi-cycle simulation trajectories.** Dynamic mass-balance model depicting the compounding feedback loop across 150 cycles (corresponding to 3 SRTs). The y-axis shows accumulated deviations (Δ accumulated change from baseline) relative to pre-co-feeding operational conditions. Asymmetric formate utilization provides an extra energy boost that drives continuous GAO glycogen and biomass accrual, enabling expanded GAO acetate uptake and the concomitant displacement of PAO biomass and acetate assimilation. Complete mathematical model formulations are provided in Supplementary Appendix File 3.

## 4. Discussion

### 4.1 Metabolic wiring and inorganic resource management shape heterotrophic niche differentiation

The findings of this study highlight how the strategic management of inorganic resources— specifically CO_2_/HCO_3_^−^ and phosphate (Pi)—shapes heterotrophic niche differentiation. Organisms facing identical macro-scale environmental conditions can evolve profoundly divergent metabolic architectures that achieve comparable ecological fitness through distinct resource routing.

Mechanistically, this divergence stems from how each guild manages intracellular redox hemeostasis. While PAOs route excess PHA-derived electrons toward aerobic respiration to power polyphosphate storage, GAOs face a massive, disproportionate carbon demand for glycogen replenishment (nearly 18-fold higher than structural biomass needs). Rather than dissipating carbon-skeletons as CO_2_, GAOs deploy heterotrophic, reductive CO_2_/HCO_3_^−^ re-assimilation via the EMC pathway. This effectively couples structural carbon conservation with tight redox control via mitigated electron overflow.

The energetic efficiency of GAOs is further fortified by a tightly synchronized suite of bioenergetic shortcuts—most notably the *PPDK–AcsA* loop halving acetate activation costs, and the *PFOR–Fpr* module generating low-potential ferredoxin. Together, these adaptations establish a highly poised redox landscape where low-potential electrons can be dynamically allocated between membrane-bound energy conservation and reductive carbon fixation as intracellular conditions demands.

Ultimately, these contrasting strategies reveal two distinct evolutionary solutions: PAOs prioritize energy storage via inorganic phosphate cycling, whereas GAOs excel in carbon conservation and bioenergetic efficiency^39^. In engineered wastewater systems, this distinction carries major operational consequences. While influent phosphate fluctuates unpredictably, inorganic carbon (CO_2_/HCO_3_^−^) is structurally guaranteed by endogenous respiratory activity. This guaranteed availability of inorganic carbon provides GAOs with a permanent ecological safety net, explaining their exceptional resilience under variable operational regimes.

### 4.2 Asymmetry in minor secondary substrate utilization drives non-linear community shifts through generational compounding

A central finding of this study is that minor secondary substrate inputs—equivalent to just 4.7% on an electron/COD basis (0.1:1 C-mol:C-mol formate-to-acetate ratio)—can exert a disproportionate, non-linear influence on long-term community assembly. This shift relies fundamentally on an asymmetric bioenergetic niche: while primary acetate is shared, minor formate co-feeding enabled *Ca.* Competibacter GAO to selectively secure an exclusive per-cycle bioenergetic advantage.

Crucially, this asymmetric advantage disrupted the cycle-boundary steady states established under sole-acetate feeding, where intracellular storage pools and biomass inventory return to baseline at the end of each operational cycle. Rather than resetting to baseline, the intracellular glycogen pools and biomass inventory of the selectively favored GAO populations systematically expanded cycle after cycle. Decoupled from hydraulic throughput, the solids retention time (SRT) did not merely waste biomass; it retained cells carrying this accumulated intracellular memory, turning a subtle per-cycle bioenergetic gain into a multi-generational ratchet that drove total population shifts over generational timescales.

These insights expose key boundary conditions that are rarely captured in established bioprocess models, such as Activated Sludge Model No. 3 (ASM3) and its Bio-P extensions^56^, which rely heavily on single-cycle kinetics, primary macronutrient stoichiometry, and cycle-boundary steady-state assumptions to forecast long-term performance. Our data demonstrate how minor, short-term metabolic advantages accumulate exponentially when SRT is decoupled from hydraulic throughput. Such iterative compounding effects are inherently masked within short-term experimental designs or steady-state models that overlook multi-cycle feedforward dynamics, underscoring the necessity of tracking cumulative intracellular polymer and biomass trajectories in multi-substrate regimes.

Consequently, resolving these compounding mechanisms is essential for translating well-controlled laboratory baselines into predictive frameworks for full-scale bioprocesses. Real-world wastewater environments rarely feature isolated substrates; instead, they are sustained by complex, dynamic carbon mixtures^57–63^. While laboratory macrocosms must simplify these streams to isolate core physiology, secondary or auxiliary substrates cannot be assumed to follow primary substrate kinetics or affect all guilds symmetrically. By systematically moving from single-substrate baselines to defined co-substrate mixtures, we can unravel these non-linear selection pressures layer by layer—ultimately transforming complex operational vulnerabilities like EBPR instability^40,64–70^ from ecological mysteries into predictable, manageable events.

### 4.3 Beyond abstraction: elevating biochemical resolution in microbial ecology

The findings of this study highlight an ongoing challenge in current microbial ecology frameworks: while functional guild classifications based on macro-scale primary substrate preferences have historically proven useful, they frequently overlook the fine-scale biochemical and stoichiometric mechanisms that ultimately dictate competitive outcomes in highly dynamic environments. As illustrated here, subtle variations in intracellular cofactor-recycling loops, ferredoxin-poised redox landscapes, and inorganic resource management can fundamentally govern competitive fitness and shape niche differentiation. The expanding availability of integrated multi-omics now makes it feasible—and necessary—to incorporate high-resolution biochemical insights into predictive frameworks of niche partitioning and community assembly across engineered and natural ecosystems. While the precise fluxes of the EMC pathway and the exact mechanics of its predicted flavin-based electron bifurcation (detailed in supporting information) await definitive biochemical purification and enzymatic characterization, our genomic architecture and proteomic alignments provide a clear blueprint for such targeted investigations.

Over two decades ago, Oremland and colleagues^71^ questioned whether environmental microbiology would wither under the weight of a purely descriptive catalog of genes, calling for a concerted coupling of metagenomic sequences with classical ecophysiological validation and biochemical isolation. Over the subsequent decades, echoing voices have continued to highlight a persistent deficit in testing fundamental ecological hypotheses or scaling laws within the multi-omics era^72^, emphasizing the urgent need to bridge high-throughput sequence data with deterministic, predictive modeling^73^. In this study, we have sought to address this enduring gap by linking macro-scale phenotypic observations with multi-omics-generated metabolic insights. While definitive biochemical validation and phenotypic isolation remain necessary next steps, we hope this work serves as an integrative case study that inspires cross-disciplinary dialogue and collaborative synergy among microbial physiologists, quantitative ecologists, biochemists, and bioprocess engineers alike.

## Acknowledgement

Mvl, TPW and YW were supported by the SIAM Gravitation Grant 024.002.002, The Netherlands Organization for Scientific Research. This study was funded by the Spinoza prize awarded to M. C. M. van Loosdrecht in 2014 by the Dutch Research Council (Nederlandse Organisatie voor Wetenschappelijk Onderzoek, NWO). YW was supported by the Delft Blue high-performance computing cluster (part of the DHPC facility) for the computational tasks in this study.

## Data Availability Statement

Raw paired-end metagenomic sequence reads and metagenome-assembled genomes (MAGs) generated in this study have been deposited in the National Center for Biotechnology Information (NCBI) database under BioProject accession number PRJNA1514754. Raw sequence reads are available in the Sequence Read Archive (SRA) under BioSample accessions SAMN62515555–SAMN62515557. The MAGs representing dominant *Ca.* Dechloromonas and *Ca.* Competibacteraceae populations are accessible in GenBank under BioSample accessions SAMN62518285– SAMN62518291. All curated proteomic signals, metabolic pathways, reaction stoichiometry (covering PHA, glycogen, anaplerosis, and ferredoxin biochemistry), and complete stoichiometric simulation models are provided in Supplementary Appendix File 2 and Supplementary Appendix File 3.

## Declaration of AI-Assisted Technologies in the Writing Process

During the preparation of this manuscript, YW used AI tools to refine the linguistic clarity and editorial formatting of the text and figure captions. Following the use of these tools, the authors reviewed, verified, and edited the final outputs to ensure absolute technical accuracy and take full responsibility for the scientific content of the work.

## Footnotes

1 Although physiological ferredoxin (E’ ≈ -450 to -520 mV) often undergoes single-electron transitions, it is treated as a two-electron carrier in the stoichiometric matrix of this study.

