## Supporting Information for "Dark CO_2_ Fixation via the Ethylmalonyl-CoA Pathway Establishes Metabolic Parity: A Stoichiometric Basis for Compounding Ecosystem Shifts"

**S1.** **Fluorescence in situ Hybridization (FISH)**

To visually corroborate the relative abundance of the dominant microbial guilds across the different experimental phases, fluorescence *in situ* hybridization (FISH) was performed according to established protocols^1^. Total bacteria were targeted using an equimolar mixture of the oligonucleotide probes EUB338 ^2^, EUB338-II, and EUB338-III probes ^3^. To target *Candidatus* Accumulibacter PAOs ^4^ and *Ca.* Dechloromonas PAOs ^5^ ], an equimolar mix of the probes Bet135, Dech443, Acc213, Acc470, Acc471, Acc471_2, Acc635, and Acc1011 was applied.. *Candidatus* Competibacter GAOs were targeted using a specific probe mixture consisting of GAOQ431 and GAOQ989^6^. Samples were mounted using Vectashield (Vector Laboratories, Burlingame, CA, USA) to preserve the fluorescent signal and counterstain total biomass. acquired using a ZEISS LSM Confocal Laser Scanning Microscope (Carl Zeiss, Jena, Germany) equipped with a 100x/1.4 Oil DIC Plan-Apochromat objective lens. Laser excitation lines and emission filters were selected according to the specific fluorophore profiles (i.e., Cy3, Cy5, FITC]), and image processing was performed using ZEISS ZEN software.

**S2. High-resolution metagenome and proteome analysis**

**S2.1 DNA extraction and metagenomic sequencing read quality control**

Biomass samples (aliquoted in 2 ml) were collected from the bioreactor during the pseudo-steady state of phase FA00 and FA02, as well as at the termination of the transient phase FA01. Samples were centrifuged at 12,000 *g* for 5 min at 4 °C to separate the biomass from the supernatant. Biomass pellets were stored at −80 °C until DNA extraction and lyophilization for protein extraction.

Genomic DNA was extracted using the DNeasy PowerSoil Pro Kit (Qiagen, Germany) according to the manufacturer’s instructions. Metagenomic sequencing was performed on an Illumina NovaSeq platform (2 × 150bp paired-end), yielding approximately 10 Gb of raw paired-end reads per sample across the three experimental phases (FA00, FA01, and FA02).

Raw paired-end sequencing reads were processed for quality control and filtered using *fastp* (v0.23.2)^7^. Adapter sequences were automatically detected and removed, duplicate reads were discarded, and poly-G tails were trimmed. Quality trimming was executed at both the 5′ and 3′ ends using a 4-bp sliding window with a minimum mean Phred quality threshold (*Q* >= 20). Reads containing more than five ambiguous (N) bases were discarded, and only reads maintaining a minimum length of 50 bp post-trimming were retained for downstream bioinformatic analysis.

**S2.2 Metagenomic assembly and co-assembly strategies**

Cleaned paired-end reads from the four individual datasets were assembled *de novo* using *metaSPAdes* (v4.2.0)^8^ with a multi-k-mer strategy (k = 21, 33, 55, 77, and 99) to maximize contiguity. Read error correction was disabled to execute only the assembly module, and default parameters were applied. To improve the recovery of low-abundance organisms, co-assemblies were simultaneously performed using *MEGAHIT* (v1.2.9)^9^ with the identical multi-k-mer strategy (k = 21, 33, 55, 77, and 99) under default settings.

**S2.3 Read mapping and coverage quantification**

To enable comparative coverage analyses, the cleaned reads from each individual sample were mapped back to both the individual assemblies and the co-assemblies using *Bowtie2* (v2.4.1)^10^ in paired-end mode. Indices were built using bowtie2-build, and the resulting alignments were output in SAM format. SAM files were subsequently converted to BAM format, sorted, and indexed using *SAMtools* (v1.3.1)^11^. Sequencing depth across the assembled contigs was calculated from the finalized BAM files via the samtools depth function. Default parameters were maintained throughout this pipeline.

**S2.4 Genome binning, dereplication, and taxonomic classification**

Automated metagenomic binning was performed on the individual assemblies and co-assemblies using *MetaBAT2* (v2.12.1)^12^ under default parameters, generating 4 distinct sets of genome bins. These bins were aggregated and dereplicated using *dRep* (v3.6.2)^13^ to yield a non-redundant set of metagenome-assembled genomes (MAGs). The completeness and contamination of the recovered MAGs were assessed via *CheckM* (v1.2.4)^14^, and taxonomic classification was assigned using the Genome Taxonomy Database Toolkit, *GTDB-Tk* (v2.5.2)^15^. Bins were evaluated based on the quantitative thresholds established by the Minimum Information about a Metagenome-Assembled Genome (MIMAG)^16^ consortium. Given our focus on metabolic pathway reconstruction, strict thresholds were maintained for completeness (>90%) and contamination (<5%)—matching the standard metrics for high-quality drafts—to ensure accurate pathway topology and prevent false-positive functional assignments for MAGs of interests. High strain heterogeneity in select abundant bins was interpreted as representative of the population's pan-genomic functional potential within the bioreactor, rather than multi-species contamination.

**S2.5 Functional genome annotation and metabolic pathway reconstruction**

Functional annotation of the MAGs was conducted using *Prokka* (v1.13)^17^ under the bacterial genetic code (translation table 11). This pipeline predicted protein-coding genes, ribosomal RNAs (rRNAs), transfer RNAs (tRNAs), and non-coding RNAs (with *Rfam* searches enabled), retaining all gene and mRNA features in the final output. Metabolic pathways and biogeochemical traits were reconstructed using *METABOLIC-G* (v4.0)^18^. Within this pipeline, protein sequences were profiled against the full KOfam database for KEGG Orthology (KO) assignment, filtering out annotations with a Hidden Markov Model (HMM) score < 0.75. All other configurations for *Prokka* and *METABOLIC-G* were kept at default values.

**S2.6 Proteome extraction and shotgun metaproteomics**

Proteins were extracted from three representative phase samples as described previously^19,20^. Briefly, approximately 6 mg of lyophilized biomass pellet was homogenized across three vortex–ice incubation cycles using glass beads (150–212 μm, Sigma Aldrich), 50 mM TEAB buffer, 1% (w/w) sodium deoxycholate and Bacterial Protein Extraction Reagent (B-PER, Thermo Scientific). Proteins in the supernatant were precipitated with 6.1 N trichloroacetic acid solution 6.1 N (Sigma Aldrich) added at 1:4 (v/v) ratio to the supernantant. The resulting pellet was washed and disrupted twice with ice-cold acetone and redissolved in 6 M urea (Sigma Aldrich). The sample was reduced with 10 mM dithiothreitol (Sigma Aldrich) at 37 °C for 60 min and alkylated with 20 mM iodoacetamide (Sigma Aldrich) in the dark for 30 min. Samples were diluted with 100 mM ammonium bicarbonate to achieve a final urea concentration below 1 M. Protein was digested overnight at 37 °C (300 rpm) using 0.1 μg/µL sequencing-grade trypsin (Promega). Solid-phase extraction was performed using an Oasis HLB 96-well elution Plate (2 mg sorbent per well, 30 *μ*m, Waters) coupled to a vacuum manifold. Columns were conditioned with MeOH, equilibrated twice with ultra-pure water, loaded with peptide samples, washed twice with 5% MeOH, and sequentially eluted with 2% formic acid in 80% MeOH and 1 mM ammonium bicarbonate in 80% MeOH. Samples were evaporated to dryness in a Concentrator Plus centrifuge (Eppendorf) at 45 °C and stored at −20 °C until analysis.

Peptide samples were reconstituted in 20 *μ*l of 3% acetonitrile and 0.01% trifluoroacetic acid, incubated at room temperature for 30 min, and thoroughly vortexed. Total protein concentration was measured at λ = 280 nm on a NanoDrop ND-1000 spectrophotometer (Thermo Scientific) and adjusted to 0.5 mg/ml. Shotgun metaproteomics was conducted as previously described ^19,20^ in randomized order with technical duplicates.

**S2.7 Metagenomic and metaproteomic-based community profile**

Community-wide protein expression dynamics across phases FA00, FA01 and FA01 were evaluated via shotgun metaproteomics, using spectral abundance as a proxy for metabolic activity and proteomic biomass allocation. The 35 high-quality (HQ) metagenome-assembled genomes (MAGs) accounted for ~80% of the total matched protein pool and 65-69% of total community DNA reads. In total, 3916 unique protein groups were detected across the 35 HQ MAGs, of which 3099 were quantified with at least two unique peptides (representing 79% of mass-normalized spectral counts).

Proteins corresponding to two PAO MAGs and six GAO MAGs were consistently detected. These eight core target MAGs accounted for ~94% of total proteomic spectral abundance and 86-87% of total metagenomic DNA reads among all 35 HQ MAGs. The PAO fraction comprised one *Ca.* Accumulibacter MAG and one *Ca.* Dechloromonas MAG (family Rhodocyclaceae), with *Ca.* Dechloromonas exhibiting overwhelming dominance. The GAO fraction was composed of one *Ca.* Contendibacter MAG and five *Ca.* Contendibacter MAG within the *Ca.* Competibacteraceae family (Fig. 2).

Metagenomic and metaproteomic relative abundances were concordant during phase FA02 (Figure 2), where the dominant GAO population exhibited both higher biovolume and numerical superiority (Figure S2). Conversely, during phase FA00 (baseline), metagenomic sequencing indicated numerical dominance of *Ca.* Dechloromonas ($>60\%$), whereas metaproteomic profiling reflected the smaller cell volume of this PAO clade, revealing that *Ca.* Dechloromonas, *Ca.* Competibacter, and *Ca.* Contendibacter actually shared comparable protein biomass allocation ($\sim30\%$ each).

Metagenomic and metaproteomic relative abundances were concordant during phase FA02 (Figure 2), where the dominant GAO population exhibited both higher biovolume and numerical superiority (Figure S2). Conversely, during phase FA00(baseline), metagenomic sequencing indicated numerical dominance of *Ca.* Dechloromonas (> 60%), whereas metaproteomic profiling reflected the smaller cell volume of this PAO clade, revealing that *Ca.* Dechloromonas, *Ca.* Competibacter, and *Ca.* Contendibacter actually shared comparable protein biomass allocation (~ 30% each).

**S3. Hypothetical anaerobic deployment of the ethylmalonyl-CoA pathway via flavin-based electron confurcation**

In addition to harvesting energy from reduced ferredoxin (Fd_red_) via the *PFOR-Fpr-PntAB* enzyme cascade described in the main text, Fd_red_ may also be partitioned to drive specific thermodynamic bottlenecks under anaerobic conditions. specifically, Fd_red_ being leveraged to power the thermodynamically challenging biochemistry. anaerobic oxidation of methylsuccinyl-CoA to mesaconyl-CoA (Reaction 2 in Figure 5) via flavin-based electron confurcation (FBEB), Specifically, the co-localization and strong expression of electron transfer flavoprotein subunits A and B (*EtfAB*) alongside methylsuccinyl-CoA dehydrogenase (*Mcd*) (Figure S7) suggests that Fd_red_ can power the endergonic anaerobic oxidation of methylsuccinyl-CoA to mesaconyl-CoA^[1]^ via flavin-based electron confurcation (FBEB; Reaction 2 in Figure 5).

While the complete, carbon-conserving ethylmalonyl-CoA (EMC) pathway is classically considered an aerobic engine, its potential anaerobic operation—linked via the *Mcd–EtfAB* complex—introduces a novel hypothesis: GAOs may couple portions of the EMC pathway with glycogen mobilization and acetate assimilation to optimize redox poising and metabolic flexibility under anaerobic conditions. Note that this anaerobic EMC model is presented strictly as a theoretical framework for future experimental validation and is omitted from our core stoichiometric matrix calculations.

**S4. Mathematical Framework and Parameters for Multi-Cycle Stoichiometric Simulations**

The stoichiometric simulations visualized in Figure 7 operate under the mass balances established in the Figure 5b and the stoichiometry detailed in Appendix file 2. The additional energy (1.24 mmol * 2.5 mol-ATP/mol = 3.09 mmol ATP) derived from aerobic formate oxidation spares a fraction (*x*) of intracellular polyhydroxyalkanoates (PHB) from energy-yielding oxidation, calculated as:

*x* = $\frac{3.09 mmol ATP}{21.5 mol ATP/mol PHB}=0.14 mmol PHB$

Reflecting the native stoichiometric routing of the GAO aerobic metabolism, this spared PHB (*x*) is dynamically split, allocating half (50%) to glycogen expansion and half (50%) to structural biomass synthesis. According to the stoichiometry that 1.6 mol of PHB is needed for the replenish of each mole of glycogen detailed in Appendix File 2, the per cycle glycogen pool expansion due to aerobic formate respiration is:

ΔGlycogen *_FA_* = $\frac{50\% * x}{1.6 mol/mol}$ = 0.313 *x* (mmol)

With the biomass yield on PHB, Y_X/PHB_ , at 0.5 C-mol Biomass/mol PHB, and the carbon content in PHB monomer at 4 C-mol/ mol-PHB, the per cycle GAO biomass expansion due to formate is:

ΔGAO-Biomass *_FA_* =4 mol/mol *50% **x* * 0.5 C-mol/mol = *x* (C-mmol)

To maintain a constant Solid Retention Time (SRT) of 10 days (at an operating frequency of 5 cycles per day; 50 cycles per SRT), 2% of this formate-induced GAO glycogen accumulated and formate-induced GAO biomass synthesized is purged via daily wasting at the end of each operational cycle, while the remaining 98% (*r* = 0.98) is retained and recycled into the subsequent anaerobic phase.

Given the theoretical anaerobic stoichiometry of a canonical GAO phenotype, the glycogen-to-acetate assimilation ratio is hardcoded at 1:2 (mol:mol, Figure 5b). Consequently, the recycled glycogen returning to the anaerobic phase of cycle *n* expands the anaerobic acetate uptake capacity of GAOs (mmol, ΔGAO-Acetate*_n_*), calculated according to:

ΔGAO-Acetate*_n_* = 2 ΔGAO-Glycogen*_n-1_*

Utilizing a theoretical biomass yield (Y*_X/S_*, Figure 6) of 0.33 C-mol Biomass / C-mol Acetate and accounting for the 2 mol carbons per mole of acetate (C_2_H_3_O_2_^-^), in cycle *n*, the extra GAO biomass synthesized during the aerobic phase from the extra acetate assimilated (ΔGAO-Biomass *_n_*_-AC_) is defined as:

ΔGAO-Biomass *_n_*_-AC_ = 0.33 * 2 mol/mol * ΔGAO-Acetate*_n_*

Extrapolating this iterative loop over *n* successive operational cycles of continuous formate co-feeding, the cumulative formate-induced glycogen accrual in GAOs (ΔGAO-Glycogen*_n_*, mmol) is mathematically resolved via a geometric series:

$$\Delta\text{GAO-Glycogen}_{n}=\sum_{k=1}^{n} \left( \Delta\text{Glycogen}_{\text{FA}}\cdot{0.98}^{k} \right)$$

Correspondingly, the cumulative formate-induced GAO biomass accrual (ΔGAO-Biomass*_n_*) remaining after sludge discharge at the end of cycle *n* is calculated as:

ΔGAO-Biomass*_n_* = (ΔGAO-Biomass *_n_*_-AC_ + ΔGAO-Biomass *_FA_* + ΔGAO-Biomass*_n-1_*) * 0.98

This stoichiometric model does not explicitly account for non-growth maintenance energies or cell decay over long-term operation, which were assumed to remain constant before and after formate co-feeding and are already included in the biomass yield coefficient; with this, we isolate and highlight the key driving force behind the compounding competitive edge of *Ca.* Competibacter GAO.

Formate-induced GAO biomass accrual (ΔGAO-Biomass*_n_*) was calculated directly from single-cycle PHB-sparing stoichiometry. Corresponding PAO biomass displacement (ΔPAO-Biomass*_n_*) was modeled based on acetate competition and 2% per-cycle SRT wastage, anchored to an assumed baseline PAO biomass:

ΔPAO-Acetate*_n_* = - ΔGAO-Acetate*_n_*

To model multi-cycle community dynamics without biasing toward specific omics modalities—given that metagenomic DNA abundance favored *Ca.* Dechloromonas PAO (~62%) while metaproteomics and quantitative FISH indicated *Ca.* Competibacteraceae GAO dominance (~67%)—the baseline reactor state (Phase FA00; Figure 2; 9.0 mmol acetate-feed/cycle, ~6.9 g biomass derived from an MLVSS content of 4.6 g/L and a 1.5 L working volume; Table 1) was parameterized assuming functional baseline parity (50% PAO - 50% GAO split; 4.5 mmol acetate-assimilation and 3.45 g biomass for each).

Under pseudo steady state, with 2% sludge discharge at the end of each anaerobic-aerobic cycle, assimilating $4.5\text{ mmol}$ acetate per cycle allows *Ca.* Dechloromonas PAO to replenish $2\%\times3.5\text{ g}=0.07\text{ g}$ PAO biomass. The PAO biomass dynamics under shrinking acetate uptake over successive cycles are calculated according to:

PAO-Biomass_n_ = (PAO-Biomass_n-1_) * 98% + (4.5 mmol + ΔPAO-Acetate_n_)* ((2%*3.5g)/4.5mmol) * 98%

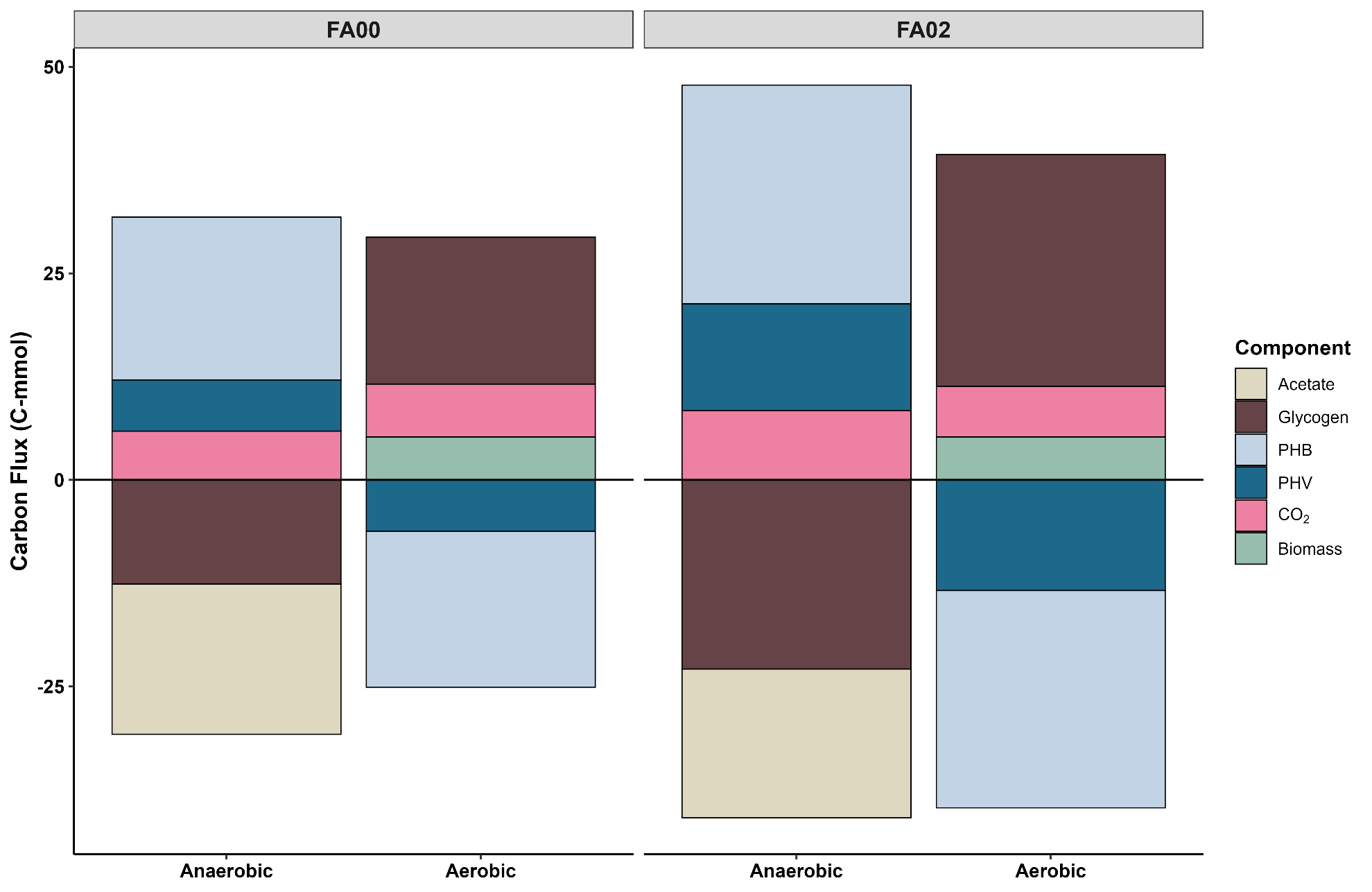

**Figure S1. Quantitative carbon flux distributions across anaerobic and aerobic cyclic phases.** Comparison of net carbon allocation (expressed in C-mmol) between the Phase FA00 and Phase FA02. Positive values denote net synthesis or accumulation of a given component, while negative values signify net consumption or degradation. Components are color-coded as detailed in the legend: Acetate (tan), Glycogen (brown), PHB (light blue), PHV (dark blue), CO_2_ (pink), and active Biomass (green). The structural shift of the community to a GAO-dominated state with EBPR deterioration is accompanied by an intensified glycogen metabolism and elevated PHV synthesis.

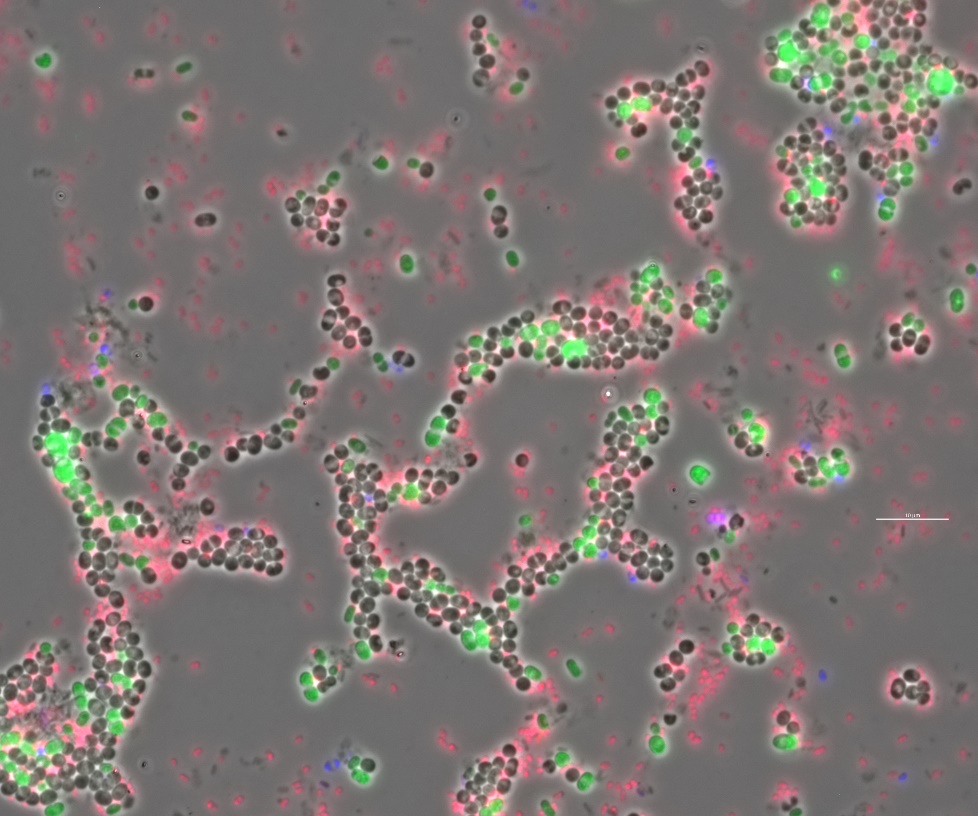

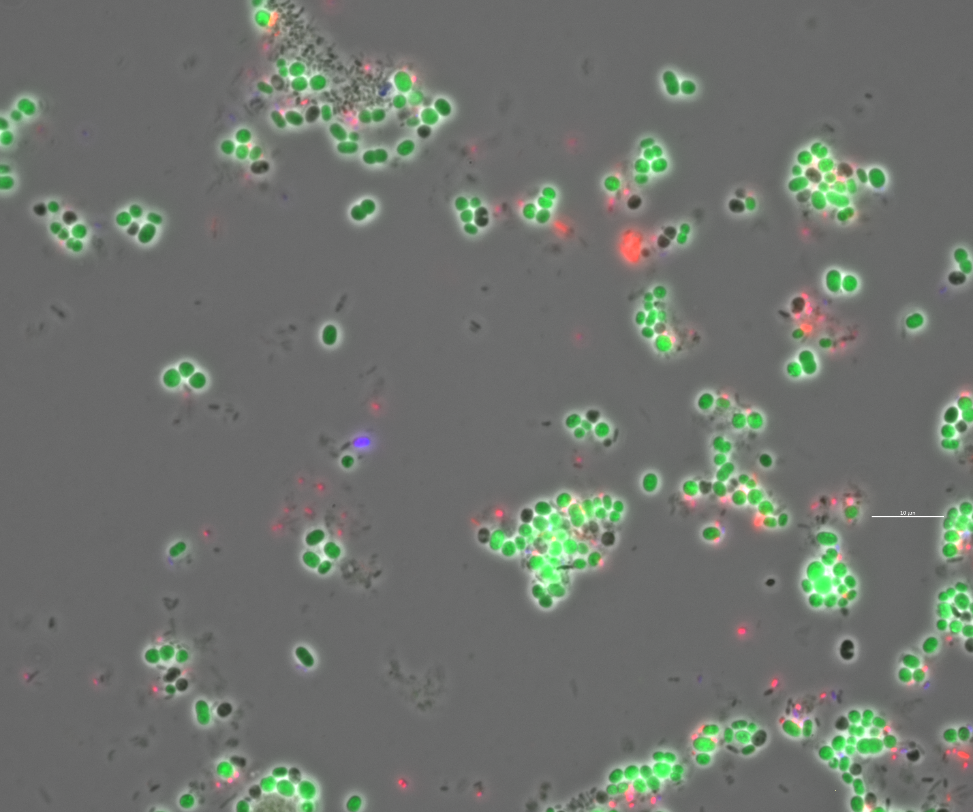

**FA02**

**FA00**

**Figure S2. Microbial community restructuring under formate co-feeding.** FISH/phase-contrast overlays showing transitions from the baseline phase (FA00) to formate co-feeding (FA02). Targets: *Ca.* Accumulibacter (blue, Cy5), *Ca.* Dechloromonas (pink, Cy3), and *Ca.* Competibacter (green, FITC). Grey cells indicate unhybridized background biomass (*Ca.* Contendobacter). Scale bar = 10 µm.

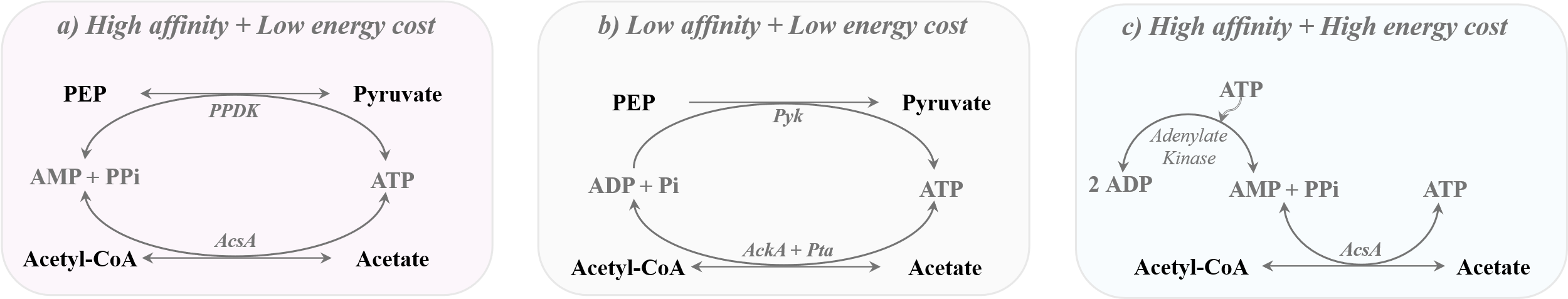

**Figure S3. Enzymatic and energetic architectures of microbial acetate activation.** Trade-offs between substrate affinity and energy cost across three activation pathways: **(a) High affinity, low energy cost:** Synchronized *PPDK–AcsA* module utilizing internal PP_i_/AMP balancing, providing a bioenergetic advantage during rapid uptake. **(b) Low affinity, low energy cost:** Classical *Pyk* and *AckA–Pta* pathway operating via standard P_i_/ADP cycling. **(c) High affinity, high energy cost:** Standalone *AcsA* coupled with adenylate kinase-mediated AMP recycling.

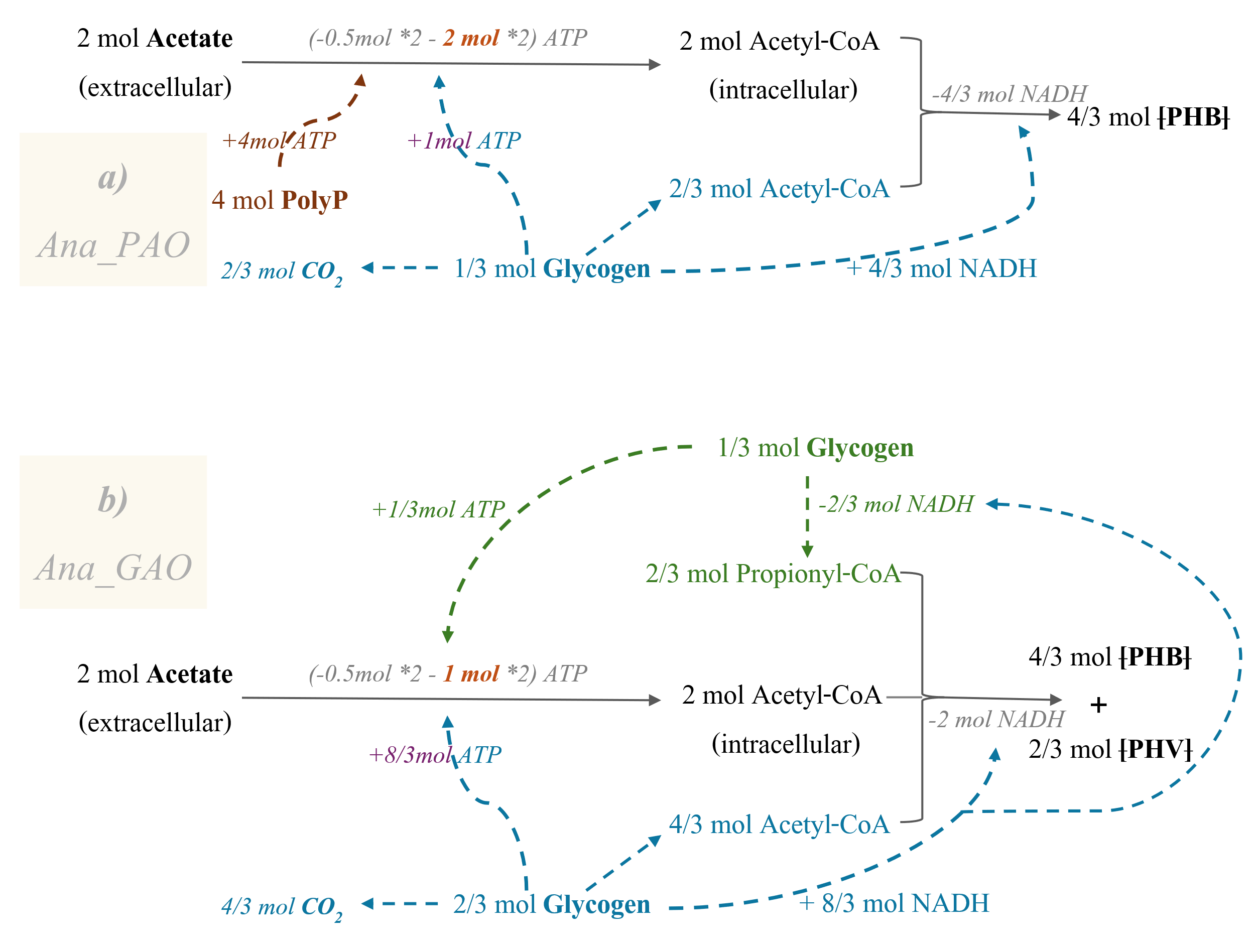

**Figure S4. Theoretical stoichiometry governing anaerobic substrate uptake and storage for PAO and GAO phenotypes. (a) Ana_PAO:** High affinity acetate activation (2 mol) via standalone *AcsA*, requiring 2 mol ATP / mol acetate from polyP cleavage alongside AMP-ADP recycling. **(b) Ana_GAO:** *PPDK-AcsA*-mediated high affinity acetate activation (1 mol ATP / mol acetate) coupled with partitioned glycogen catabolism via glycolysis (teal) and the reductive TCA branch (green) to balance energy and redox (NADH) demands. The model accounts for glycolytic ATP yields, where ferredoxin-mediated pyruvate oxidation yields 4 mol ATP / mol glycogen in GAOs versus 3 mol ATP in PAOs. (Assumed acetate transport cost = 0.5 mol ATP / mol acetate).

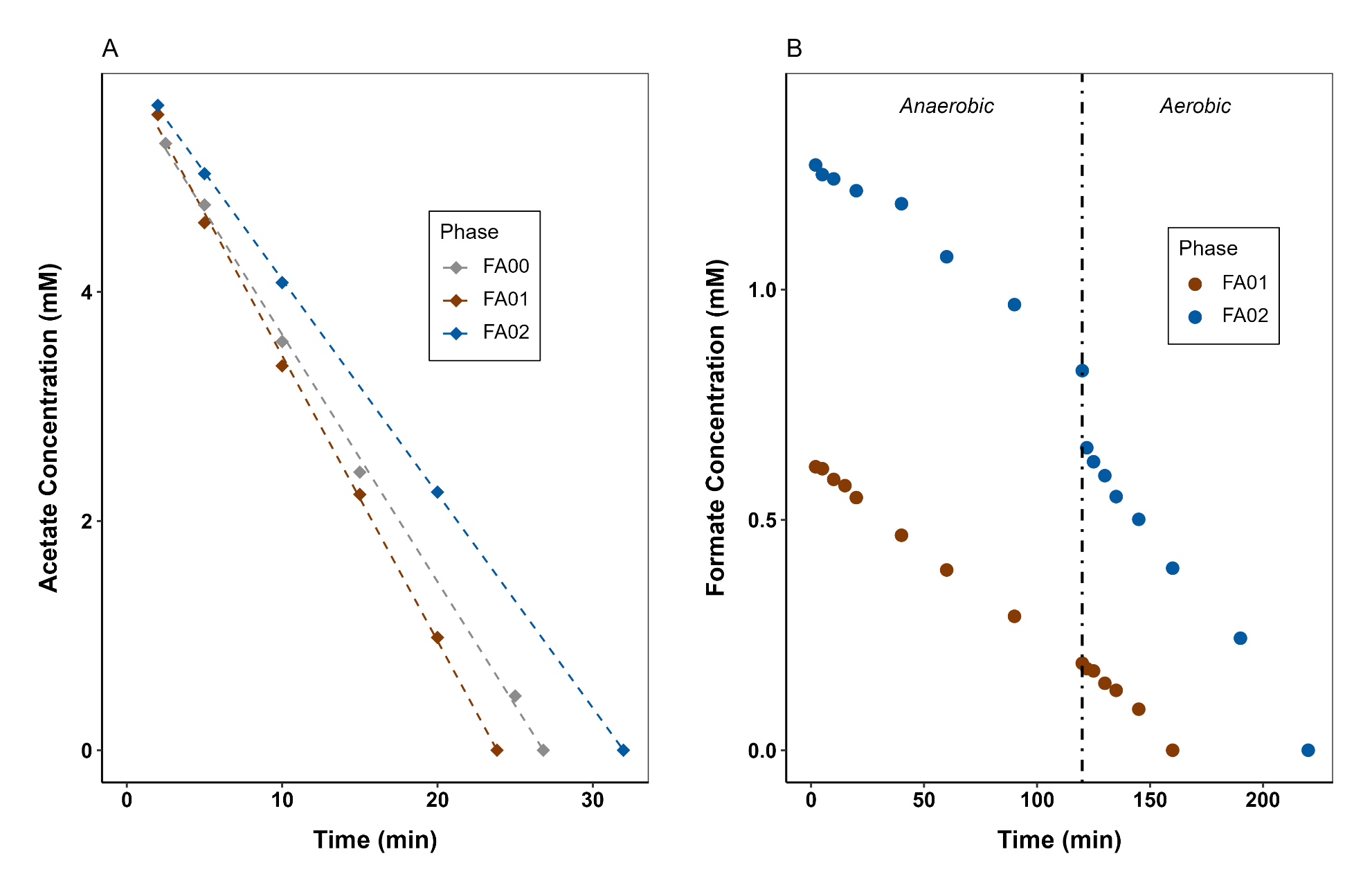

**Figure S5. Substrate consumption kinetics across operational phases. (A)** Anaerobic acetate depletion profiles in the baseline phase (FA00), transient phase (FA01), and steady-state co-feeding phase (FA02). **(B)** Formate consumption dynamics across the cycle. The vertical dash-dotted line marks the anaerobic–aerobic transition (*t* = 120 min). Under steady-state co-feeding (FA02), approximately 69% of the initial formate pool carries over into the aerobic phase.

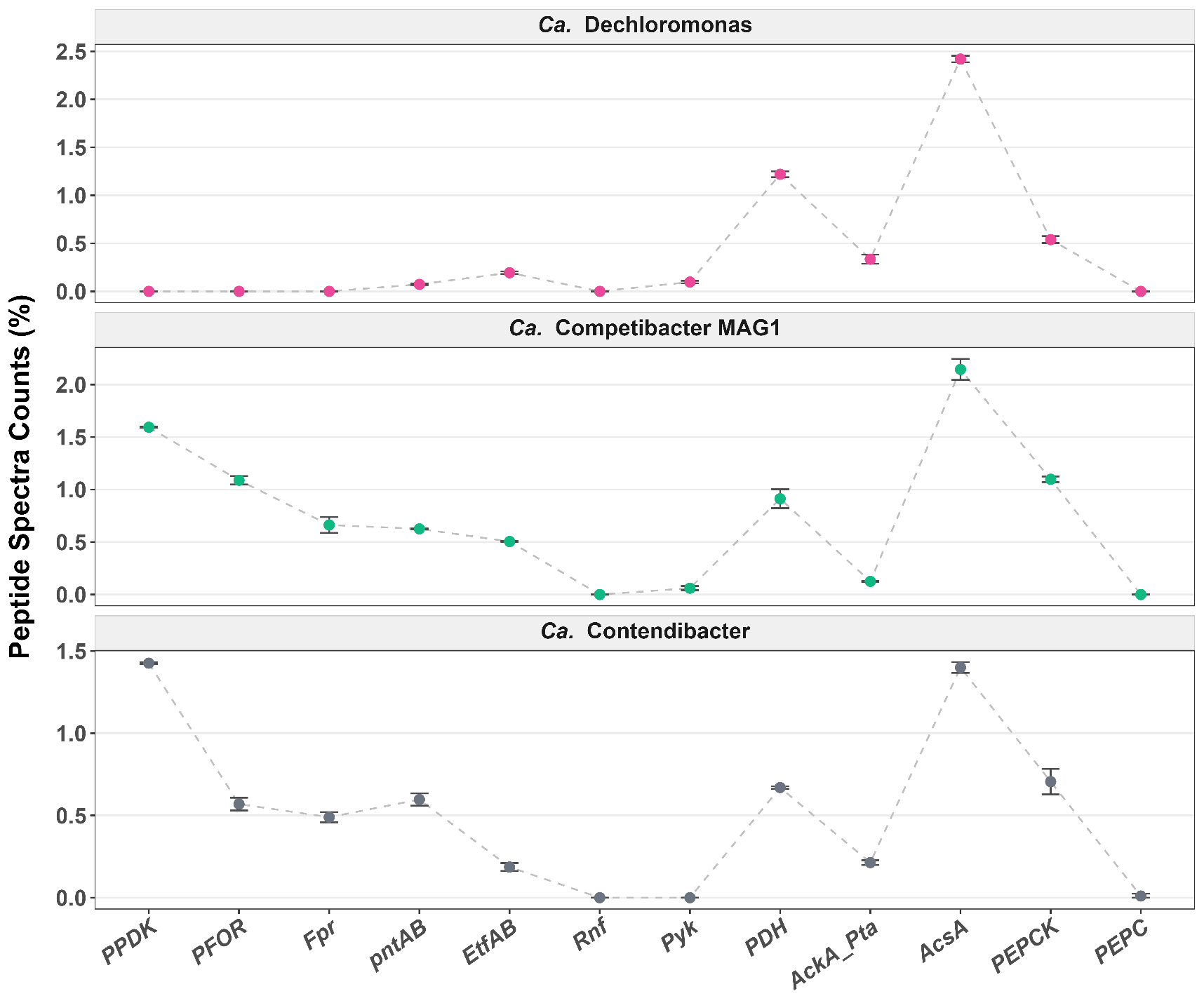

**Figure S6. Metaproteomic expression profiles of central metabolic nodes across dominant enrichment lineages.** Relative abundance of core enzymes when each target population reached its peak community abundance: *Ca.* Dechloromonas (top; FA00), *Ca.* Competibacter MAG1 (middle; FA02), and *Ca.* Contendibacter (bottom; FA00). Values represent the percentage of total peptide spectral counts mapped per genome (mean ± SD, *n*=2; note independent y-axis scales). GAO-specific upregulation: GAOs selectively express pyruvate phosphate dikinase (*PPDK*), pyruvate:ferredoxin oxidoreductase (*PFOR*), ferredoxin-NADP⁺ reductase (*Fpr*), transhydrogenase (*PntAB*), and electron transfer flavoprotein (*EtfAB*). PAO-specific preference: *Ca.* Dechloromonas PAO relies primarily on classical pyruvate dehydrogenase (*PDH*) and pyruvate kinase (*Pyk*). Shared architecture: All lineages rely on high-affinity acetyl-CoA synthetase (*AcsA*) over low-affinity acetate kinase/phosphotransacetylase (*AckA–Pta*) system, and reversible phosphoenolpyruvate carboxykinase (*PEPCK*) over phosphoenolpyruvate carboxylase (*PEPC*). Undetected complexes: Ferredoxin:NAD^+^ oxidoreductase (*Rnf*) expression remained below detection limits across all lineages. Biochemical reactions of each enzyme are summarized in Table S2.

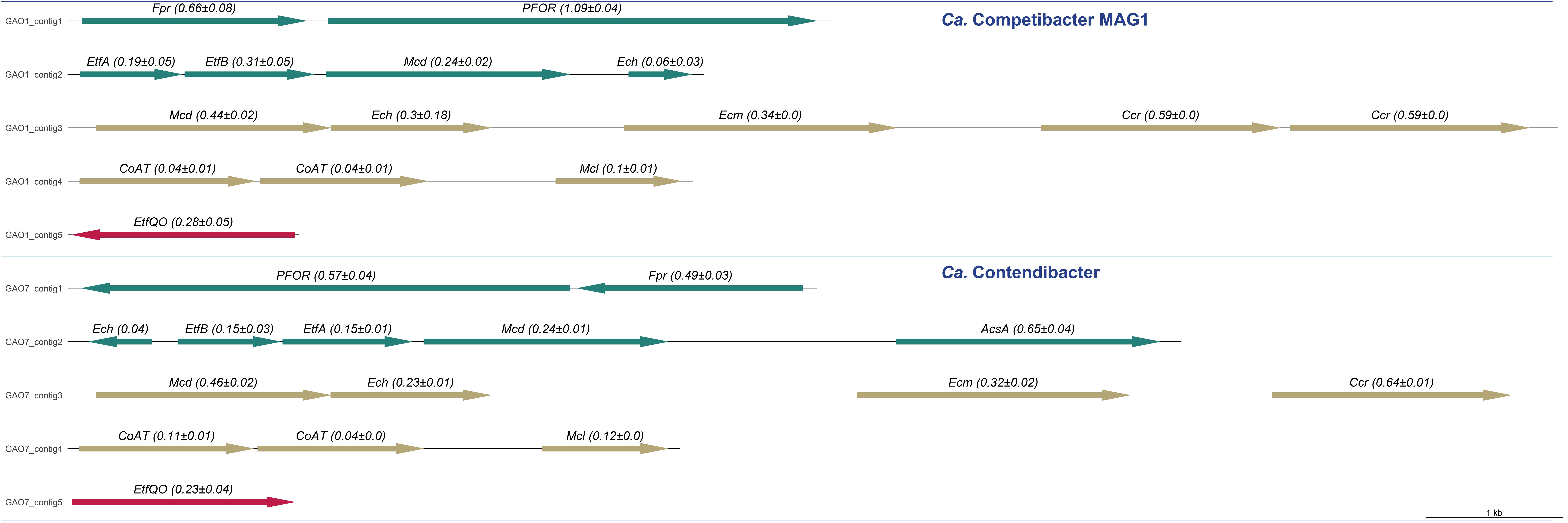

**Figure S7. Genomic architecture and metabolic operons of the ethylmalonyl-CoA pathway.** Synteny of core catalytic elements (gold), alongside anaerobic (dark green) and aerobic (red) redox management components in *Ca.* Competibacter MAG1 (upper panels) and *Ca.* Contendibacter (lower panels). Arrow direction indicates transcriptional orientation. Values in parentheses denote relative abundance quantified within the respective metaproteome (mean percentage of total spectral counts ± standard deviation). Scale bar = 1 kb. Core catalytic genes (Gold): *Mcd*, methylsuccinyl-CoA dehydrogenase; *Ech*, ethylmalonyl-CoA/mesaconyl-CoA hydratase; *Ecm*, ethylmalonyl-CoA mutase; *Ccr*, crotonyl-CoA reductase; *CoAT*, acetoacetyl-CoA:acetate CoA-transferase; *Mcl*, malyl-CoA lyase. Anaerobic redox components (Dark Green): *Fpr*, ferredoxin-$\text{NADP}^{+}$ reductase; *PFOR*, pyruvate:ferredoxin oxidoreductase; *EtfA/EtfB*, electron transfer flavoprotein subunits A and B; *AcsA*, acetyl-CoA synthetase. Aerobic redox components (Red): *EtfQO*, electron transfer flavoprotein-ubiquinone oxidoreductase.

**Table S1. Biochemistry of key enzymes involved in formate utilization and hydrogen metabolism.** Both formate hydrogenlyase (*FHL*-2) and NAD-reducing hydrogenase (*HoxHYF*) show absent or negligible expression in *Ca.* Dechloromonas. Electrons derived from H₂ oxidation by membrane-bound respiratory uptake hydrogenase (*Hyd*-2) transfer to the membranous quinone pool (QH₂), yielding approximately 0.5 mol ATP equivalents via proton motive force (PMF). Both *Hyd-4* (within the *FHL-2* complex) and *Hyd-2* are selenium-dependent [NiFeSe]-type hydrogenases, providing elevated catalytic rates and oxygen tolerance well-suited to cyclic anaerobic–aerobic ecosystems. Population-specific expression trajectories are shown in Figure 3.

| **Abbreviation** | **Gene name** | **Cellular component** | **Reaction** |
| --- | --- | --- | --- |
| *FHL*-2 (*Hyf*-NiFeSe *Hyd4*) | Formate hydrogenlyase | Membrane-bound | Formate = H_2_ + CO_2_ + PMF (~0.5 ATP) |
| *HoxHYF* | NAD-reducing hydrogenase | Cytosolic | H_2_ + NAD^+^ = NADH + H^+^ |
| *Hyd-*2 (NiFeSe H_2_ase) | Respiratory H_2_-uptake hydrogenase | Membrane-bound | H_2_ + Q = QH_2_ + PMF (~0.5 ATP) |

**Table S2. Gene copy numbers and corresponding biochemical reactions governing anaerobic acetate activation and phosphoenolpyruvate/pyruvate core metabolism across dominant lineages.** Population-specific expression trajectories for these translated gene products are visualized in Figure S6.

| **Abbreviation** | **Reaction** | **Gene Copy Number in** | | |
| --- | --- | --- | --- | --- |
|  |  | *Ca.* Dechloromonas | Ca. Competibacter | Ca. Contendibacter |
| *PPDK* | ATP + Pyruvate + Phosphate = AMP + PEP + Diphosphate | 0 | 1 | 1 |
| *Pyk* | PEP + ADP → Pyruvate + ATP | 1 | 1 | 0 |
| *AckA* | ATP + Acetate = ADP + Acetyl phosphate | 1 | 1 | 1 |
| *Pta* | Acetyl phosphate + CoA = Acetyl-CoA + Pi | 1 | 1 | 1 |
| *AcsA* | ATP + Acetate + CoA = AMP + Diphosphate + Acetyl-CoA | 1 | 1 | 1 |
| *PFOR* | Pyruvate + CoA + Fd_ox_ === Acetyl−CoA + CO_2_ + Fd_red_ | 0 | 1 | 1 |
| *Fpr* | Fd_red_ + NADP^+^ === Fd_ox_ + NADPH + H^+^ | 1 | 1 | 1 |
| *RnfABCDEFG* | Fd_red_ + NAD^+^ === Fd_ox_ + NADH + H^+^ + PMF (~0.5 ATP) | 0 | 1 | 1 |
| *PDH* | Pyruvate + CoA + NAD^+^ → Acetyl−CoA + CO_2_ + NADH + H^+^ | 1 | 1 | 1 |
| *pntAB* | NADH + NADP^+^ + PMF (~ 0.5 ATP) === NADPH + NAD^+^ | 1 | 1 | 1 |
| *EtfAB_Mcd* | Fd_red_ + NAD^+^ + Methylsuccinyl-CoA ==Mesaconyl-CoA+ Fd_ox_ + 2NADH + 2H^+^ | 0 | 1 | 1 |
| *PEPCK* | Oxaloacetate + GTP ==== Phosphoenolpyruvate + GDP + CO_2_ | 1 | 1 | 1 |
| *PEPC* | Oxaloacetate + Pi ===> Phosphoenolpyruvate + CO_2_ | 0 | 1 | 1 |

**Table S3. Stoichiometric estimation of anaerobic/aerobic ATP yields and conserved carbon capital from formate co-feeding.** ΔATP metrics define energy gained via respiratory consumption of formate (ΔFormate), which spares PHB from energy catabolism in GAOs and routes saved carbon toward GAO glycogen accumulation (ΔGAO Glycogen) and GAO biomass synthesis (ΔGAO Biomass-C) in the stable *Ca.* Competibacter-dominated phase (FA02).

| mmol | ΔAcetate /Cycle | ΔFormate  /Cycle | | ΔFormate  /Ana | ΔFormate  /Ae | ΔATP  /Ana_FA_ | ΔATP  / Ae_FA_ | PHB Spared  (aerobic) | | ΔGAO Glycogen | ΔGAO Biomass-C | |
| --- | --- | --- | --- | --- | --- | --- | --- | --- | --- | --- | --- | --- |
| FA00 | 9 | | 0 | 0 | 0 | 0 | 0 | 0 | 0.00 | | | 0.00 |
| FA02 | 9 | | 1.8 | 0.56 | 1.24 | 0.28 | 3.1 | 0.14 | 0.04 | | | 0.14 |

1. Mechanistically, while the standard reduction potential of the free methylsuccinyl-CoA/mesaconyl-CoA redox couple is approximately -125 mV, , the midpoint potential of the enzyme-bound FAD cofactor in acyl-CoA dehydrogenases (specifically *Mcd*) typically shifts to approximately -40 mV upon substrate binding.
